# A Computational Model of Stem Cell Molecular Mechanism to Maintain Tissue Homeostasis

**DOI:** 10.1101/2020.03.02.972968

**Authors:** Najme Khorasani, Mehdi Sadeghi, Abbas Nowzari-Dalini

## Abstract

Stem cells, with their capacity to self-renew and to differentiate to more specialized cell types, play a key role to maintain homeostasis in adult tissues. To investigate how, in the dynamic stochastic environment of a tissue, non-genetic diversity and the precise balance between proliferation and differentiation are achieved, it is necessary to understand the molecular mechanisms of the stem cells in decision making process. By focusing on the impact of stochasticity, we proposed a computational model describing the regulatory circuitry as a tri-stable dynamical system to reveal the mechanism which orchestrate this balance. Our model explains how the distribution of noise in genes, linked to the cell regulatory networks, controls cell decision-making to maintain homeostatic state. The noise control over tissue homeostasis is achieved by regulating the probability of differentiation and self-renewal through symmetric and/or asymmetric cell divisions. Our model reveals, when mutations due to the replication of DNA in stem cell division, are inevitable, how mutations contribute to either aging gradually or the development of cancer in a short period of time. Furthermore, our model sheds some light on the impact of more complex regulatory networks on the system robustness against perturbations.

## 1. INTRODUCTION

Throughout development, stem cells play a key role during multiple morphogenetic processes, such as tissue growth, regeneration, and repair. Stem cells are characterized by their capacity to self-renew and to differentiate to more specialized cell types [1, 2] and a balance between these two processes is necessary to maintain homeostasis in adult tissues [3–6]. Abnormalities in the differentiation or imbalance between proliferation rate and tissue demand can lead to dysfunctional tissues or tumorigenesis. On the other hand, to develop a tissue with hundreds of different cell types from a single stem cell, a non-genetic diversifying mechanism is required. Hence, understanding the underlying mechanisms which regulate the non-genetic diversity and orchestrate the stem cell proliferation/differentiation balance in the dynamic stochastic environment of a tissue is a central challenge in adult stem cell biology [7].

Stochasticity is an inevitable part of most cellular processes (including cell division) and arises from a plenitude of sources such as variation in gene expression, metabolic activities, protein and RNA degradation, etc [8–11]. This stochsticity, also called intrinsic noise, which results from the probabilistic nature of any biochemical system with a low number of reacting molecules, can lead to cell-to-cell variability during development [11]. Despite the presence of noise, a precise and robust regulation of key reactions in the cell is required for survival and functionality. An increasing number of theoretical and experimental studies are aimed at unraveling the importance of noise in such robust biological processes [11–13]. It is known that biological systems can utilize and regulate this stochasticity to improve their fitness via phenotypic variations [14, 15] and population heterogeneity [10, 11, 16, 17]. During division process, a stem cell utilizes a stochastic cell-fate decision making process, to divide either symmetrically to two differentiated (DD-division) or two new stem cells (SS-division), or asymmetrically to one differentiated and one stem cell (SD-division) [18, 19]. In an adult tissue,in homeostasis state, a perturbation leading to a dominant rate of any of the symmetric division types causes imbalance between proliferation and differentiation, which consequently diminishes the phenotypic diversity. Therefore, a robustly regulated stochastic decision-making process enhances morphogenetic processes by maintaining both proliferation/differentiation balance to avoid tissue depletion or abnormal growth [2, 5, 6] and a non-genetic diversity which is critical to the survival of living systems in noisy environments [20–25]. Cellular regulatory networks are known to play a crucial role in adjusting the decision-making mechanism by considering the effects from permanent intrinsic noise associated with living cells. Such regulatory networks have been studied extensively in a variety of organisms spanning from viruses to mammals [20]. These networks are known to control decision making from viruses [26–28] to bacteria [29–32], yeast [33] and human embryonic stem cells [34–38].

Taking into account the noisy dynamics of a small number of contributing determinants associated with intracellular processes, it is necessary to utilize a stochastic model to gain a better understanding of the behaviour of such regulatory networks. In this model, the system state is described as quantized fractions of full capacity of each determinant and can evolve stochastically over time [39]. Therefore, the probability of the system being in a given state changes with time, and cell character cannot be predicted deterministically as it is influenced by the intrinsic noise [11, 39]. To simulate the time evolution, it is suitable to use Gillespie algorithm which is proven effective for describing the trajectory of systems including a small number of determinants driven by inherent fluctuations [11, 40]. Averaging over enough simulation runs can provide us with an asymptotic approximation to the exact numerical solution of the master equation without having to deal with intractable mathematical solutions.

By focusing on the impact of stochasticity during cell-fate decision-making process, here, we propose a computational model to reveal the mechanism which regulates the proliferation/differentiation balance in a hypothetical adult tissue. In the most simple model, it is assumed that a developing tissue, consisting of stem cells and two differentiated cell types, has the tendency to maintain a homeostatic state. The proposed model is defined based on five material principles which has been discussed in [10] to study biofilm formation and they are reconsidered as follows i) stochasticity due to an intrinsic noise is a fundamental part of any living cell [10, 17, 41–46]. ii) the non-deterministic position of the cell division plane and nonuniform distribution of determinants in the cell imply that the cytoplasmic molecules are distributed randomly among daughter cells during cell division [10, 47–53]. iii) determination of cell fate by an internal switch upon the completion of cell division [20, 54]. Cell fate is assumed fixed during cell life cycle [10]. iv) the decision bias in the internal switch is determined by model parameters representing interactions between the switch elements [10]. v) a switch with more contributing components would be more stable against environmental fluctuations [10, 36].

Inspired by previous studies that revealed the impact of regulatory networks on the stability of biological systems [11, 14, 27, 33, 41, 55–58], here, we introduce a tristable switch described by a set of ordinary differential equations (ODEs) which is a formal framework to study the regulatory circuitries [36, 59]. The evolution of our inherently stochastic system is simulated by the Gillespie algorithm.

The overall outcome of our model implies that the presence of controlled noise in a population of genetically similar cells with the same environmental condition is necessary to develop population heterogeneity and also homeostasis. Furthermore, by changing the parameters in cell regulatory switches, we investigate cellular decision-making bias emanating from the stochastic environmental factors. We show that, by having enough information about the noise, predicting the cell fate after cell division is possible and that, the offspring inherit these information. Finally, to further illustrate how a transition from homeostasis to tissue depletion or abnormal growth occur in our model, we explore the behaviour of the populations consisting of cells with mutated internal switches. We show that the switches with more contributing elements are more robust against mutations. Although mutations in the stem cell usually triggers differentiation and consequently rapid depletion of stem cell population over time, accumulation of mutations leads to rapid proliferation of stem cells which is a potential indication of cancer initiation.

## 2. MATERIALS AND METHODS

### A. Cell growth and division in the population

To study the regulatory mechanism which provides the proliferation/differentiation balance in homeostatic state, we proposed a computational model described by a set of ordinary differential equations (ODEs) which was previously used in several studies to model the regulatory circuitries as tri-stable dynamical systems [36, 59]. The following set of ODEs are employed to describe a two-element regulatory switch in our model:

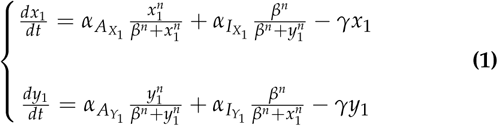

In this model, It is assumed that the cell type is controlled by the relative amount of two cytoplasmic cell fate determinants, namely *X*_1_ and *Y*_1_ whose interactions can be described in a form of a tri-stable regulatory switch (see Figure 1.a). The dynamical behavior of the determinants *X*_1_ and *Y*_1_ is studied by considering their mutual repression and self-activation effects which are modeled in the form of a Hill function [10, 27], and their degradation rate. Here, *n* is the Hill coefficient, *β* is the effective rate of determinnats synthesis, 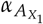 and 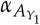 are self-activation rates, 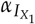 and 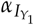 are inhibition rates, and gamma is the degradation rate.

**Fig. 1.**
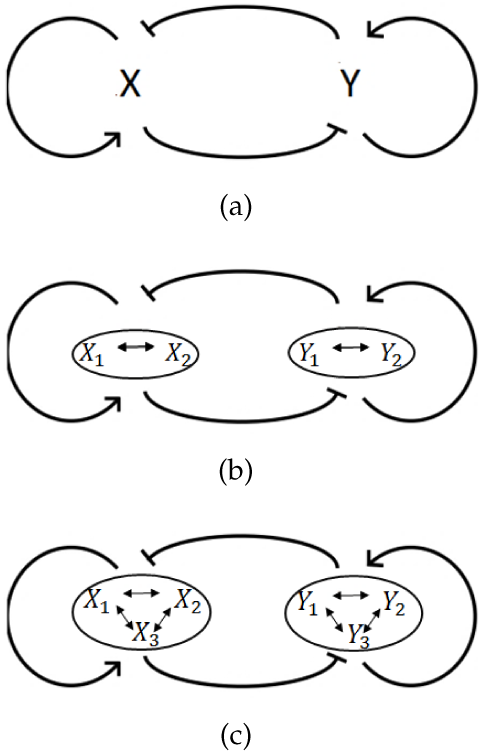
Tri-stable regulatory networks. (a) Two-element switch. (b) Four-element switch. (c) Six-element switch.

Figure 2 illustrates the described system dynamics which is visualized in a force-field representation. The grid point dimensions represent the number of determinants *X*_1_ and *Y*_1_ and each arrow shows the most probable direction which the number of determinants tends to be updated to, in each time step and based on Equation 1. The longer arrows represent a cell state with the higher tendency to fluctuate, meanwhile the more stable states of a cell is represented by shorter arrows.

**Fig. 2.**
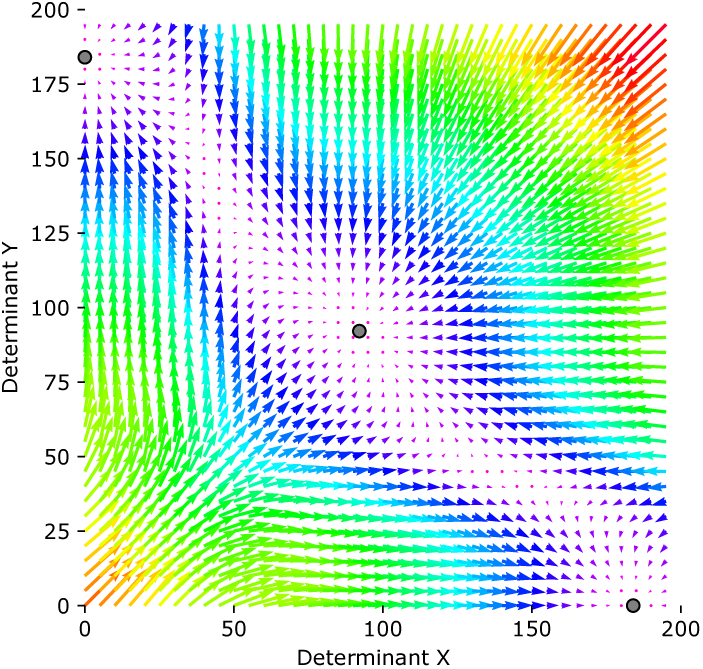
Force-field representation of the tri-stable dynamical system of a regulatory switch consisting of two cell fate determinants,namely *X*_1_ and *Y*_1_, with self-activation and mutual-repression interactions

Figure 3 shows the Waddington’s epigenetic landscape which was first described in [60]. It is derived from Equation 1, using the algorithm which is poroposed in [61]. It governs the dynamic behavior of the regulatory switch of our model. The Waddington’s landscape portrays branching ridges and valleys which represent the either-or situations which a dividing cell deal with. The cell decisions lead to one of the attractors of the regulatory switch which determines the cell final fate. When a daughter cell is born, it can be represented by a point on the surface, as the quantitative view of the cell, of Figure 3. The coordinates of the point demonstrate the value of determinants *X*_1_ and *Y*_1_ in the new born cell and determines which path should be followed to reach the final fate (one of the three attractors).

**Fig. 3.**
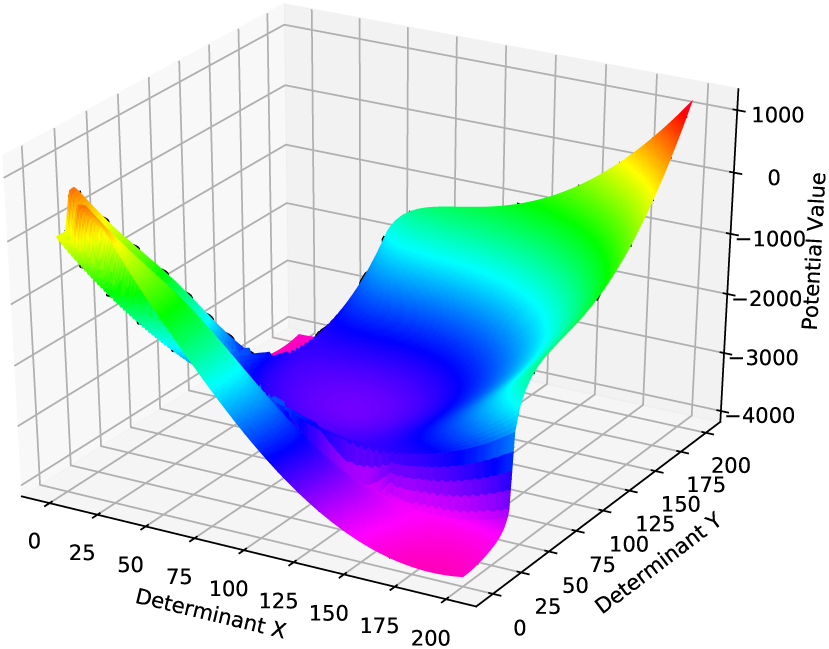
Potential landscape representation of the tri-stable dynamical system of a regulatory switch consisting of two cell fate determinants,namely X and Y, with self-activation and mutual-repression interactions

The parameters of Equation 1 are set in such a way that there would be three stable steady states, as it is shown in Figure 2, corresponding to three different cell fates, stem cell type *C* (middle attractor) and differentiated cell types *A* (bottom right attarctor) and *B* (top left attarctor). The number of determinants of *X*_1_ (*Y*_1_) involved in attractor *A* (*B*) is much larger than those of *Y*_1_ (*X*_1_). In attractor *C*, however, both determinants *X*_1_ and *Y*_1_ are involved in balance. Figure 4 represents the domains of the three attractors, *A, B*, and *C*, with three different colors, green, orange, and yellow, respectively. Each daughter cell with specific value of *X*_1_ and *Y*_1_, right after birth, can be shown as a point in Figure 4. The value of *X*_1_ and *Y*_1_ determines which attractor the cell would be absorbed to, and based on that it defines the domains of three attractors. In other word, each cell fate can be determined and fixed exactly after division based on the number of determinants *X*_1_ and *Y*_1_ in the daughter cell.

**Fig. 4.**
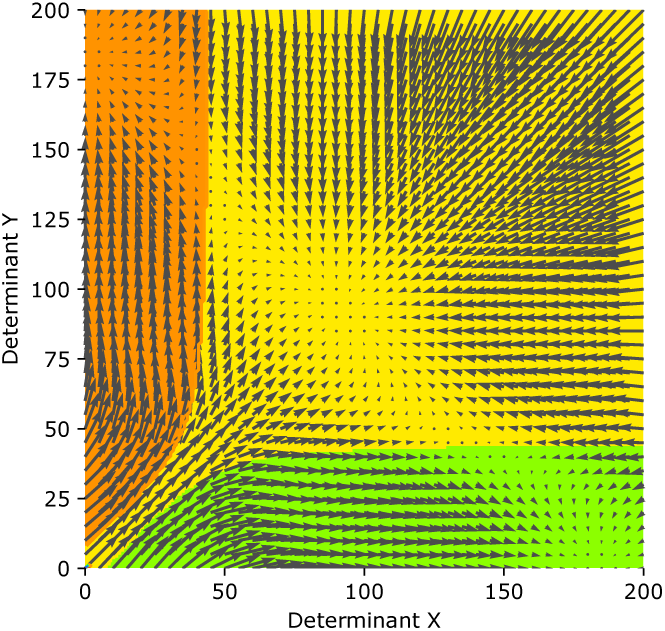
Three attractors domains corresponding to the initial values of determinants in a daughter cell.

The determinant fluctuations are captured by the Gillespie algorithm [10, 40] which is known as the gold standard for simulating models whose stochasticty arises from the small discrete number of reactant molecules[62]. In each time step, two main processes can occur, cell division and the cell determinants interactions. Therefore, five different reactions can potentially happen in each step, division and increasing/decresing of *X*_1_ value, increasing/decreasing of *Y*_1_ value. In each iteration, one of the above-mentioned processes occurs, time is updated, and the simulation continues for a whole cell cycle time *T*, where *T* = *log*(*N*) and *N* is the maximum number of cell determinants in the steady state. Hence, one can be sure that each cell can reach an attarctor in this period and it can not easily get out of that [63].

As it is shown in Equations 2, 3, 4, and 5, according to the Gilespie algorithm, one probability is assigned to each reaction. In these equations, *P*(*⋕X*_1_ → *⋕X*_1_ + 1), *P*(*⋕X*_1_ → *⋕X*_1_ − 1), *P*(*⋕Y*_1_ → *⋕Y*_1_ + 1), and *P*(*⋕Y*_1_ → *⋕Y*_1_ − 1) are the probabilities assinged to the increasing/decresing of *X*_1_ value, and the increasing/decreasing of *Y*_1_ value, respectively. As it is shown in Equations 6, 7, 8, 9, and 10 the probabilities of the four above-mentioned reactions are chosen based on the corresponding terms in Equations 1. The division probability would be equal to 1/*T*, so each cell divides at least once during the simulation.

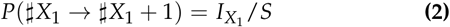

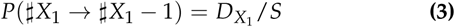

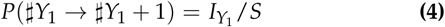

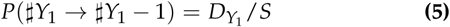

where,

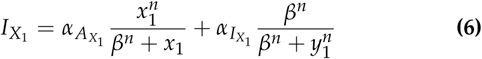

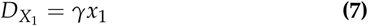

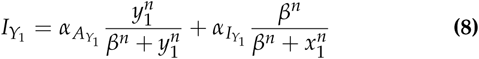

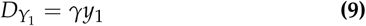

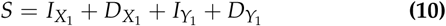

The simulation starts with a population of 50 stem cells, and the number of determinants *X*_1_ and *Y*_1_ are initialized randomly from the middle attractor region (Figure 2). As the number of determinants in the cell are updating, their corresponding trajectory in the phase plane is changing and finally reaches the domain of their attractor. As mentioned before, for each cell four reactions can happen, as a result at each time step, 4 × *⋕cells* = 4 × 50 = 200 updating reactions or one division process can occur.

Because of randomly distribution of mother cell cytoplasmic molecules between daughter cells and the non-deterministic position of the division plane [10, 47–53], here the number of determinants in each daughter cell is assumed to be from binomial distribution [64] with parameters specified according to the whole number of determinants in the mother cell, and probability of success for each trial, 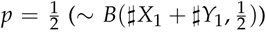. At the time of birth, each offspring phenotype is determined based on the number of determinants, which is corresponding to a coordinate in the three-region phase plane which is demonstrated in Figure 4.

Based on the off-springs fate right after division, there are two types of cell division, symmetric and asymmetric (Figure 5). The symmetric division leads to the birth of two stem cells (SS division) or two differentiated cells (DD division), while the asymmetric division generates one stem cell and one differentiated cell (SD division) [19]. In other word, stochastic partitioning of cytoplasm during cell division and the random distribution of molecules in the cytoplasm determines the division types which play a key role in maintaining the proliferation/differentiation balance in homeostatic state.

**Fig. 5.**
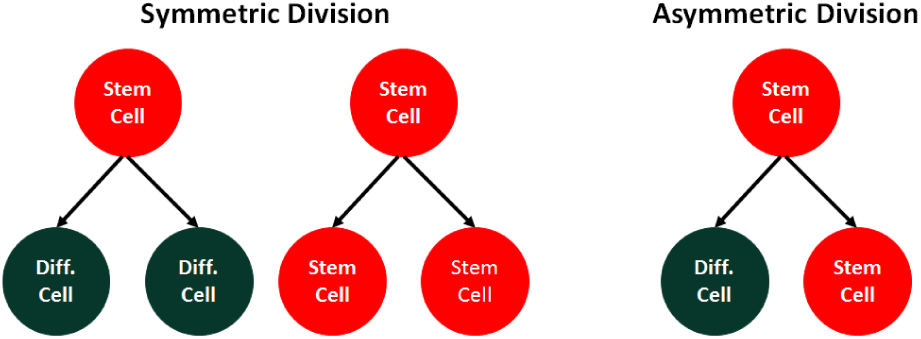
Two different types of divisions.

### B. More complex switches

It is assumed that the interactions between two determinants *X*_1_ and *Y*_1_ determine the cell fate right after cell division. The dynamics of the system is described as it is presented in Equation 1. This type of regulatory switches (Figure 1.a) are so sensitive to mutations and perturbations that directly affect the cell fate in the population. It suggests designing more complex regulatory networks consisting of a pair of clusters, with multiple elements in each, to determine the final cell fate[36]. It is revealed that these hypothesized clusters existed in biological regulatory circuitries through a literature review [65–78].

The extended switch results in more robustness against perturbations. The buffering effect is achieved by presence of more elements and the positive feed-backs in each cluster[36]. It is expected that this effect would be even stronger in more complex switches, which is in agreement with the Waddington’s idea of “*canalisation*” in [79]: “*canalisations are more likely to appear when there are many cross links between the various processes, that is to say when the rate of change of any one variable is affected by the concentrations of many of the other variables*”.

In our proposed model, each element in a cluster can have a master or supportive role in cell fate decision making. This model is in contrast with the computational model studied in [36], where all elements of the same group have identical effects in determining cell final fate. In our extended regulatory switch (Figures 1.b, 1.c), it is supposed that there is a master cell fate indicator in each cluster, and that all other elements support and regulate its effects. In other words, the different elements of the same cluster have different effects on final cell fate, which is supported by experimental observations[38].

To check the robustness of the extended model, two other ODE systems are designed in Equations 11 and 12. In Equation 11 (Equation 12) it is assumed that there are two clusters involving in cell fate decision making where they interact with each other in a four-element switch (six-element switch). Besides, clearly the elements in the extended switches can be divided in two groups (*x*−group and *y*−group). Determinants in the same group activate each other while they have a negative mutual interaction with the opposite group components.

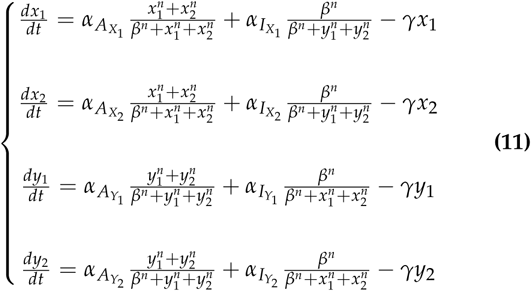

The four- and six-element switches resemble the former switch in representing a tri-stable system. The number of all elements in the *x*−group (*y*−group) involved in attractor *A* (*B*) is much larger than those in the *y*−group (*x*−group). In attractor *C*, all the elements of both groups are involved in balance. However, the elements of *X*_1_ and *Y*_1_ are the master indicators which determine the daughter cell fate after division. As it was mentioned before, all other elements in the same cluster (of the switch) only play a key role in buffering the perturbation effects on master determinants. As it is impossible to represent four/six-dimensional plots, the corresponding phase planes of four/six-element switches are plotted in two-dimensional plane. Therefore, both phase planes resemble the one of the two-element switch in Figure 2, presenting only *x*_1_ and *y*_1_ on *x* and *y* axis, respectively.

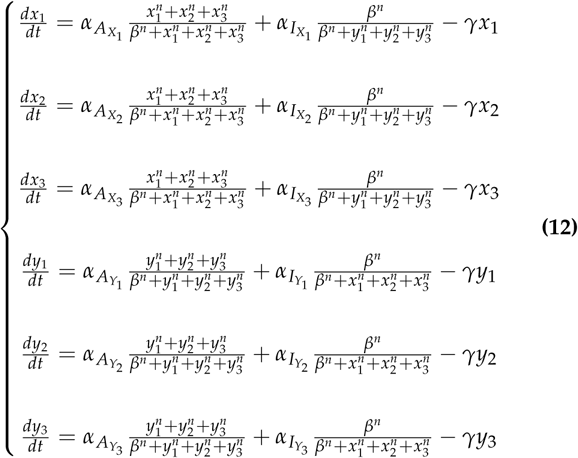

## 3. RESULTS

### A. The homeostatic state in our model

In the homeostatic state, a balance between two processes of differentiation and proliferation is necessary to maintain a fixed number of stem cells in an adult tissue [4, 5]. In our model the parameters in Equations 1, 11, and 12 are set to the values provided in Table 1. A tri-stable system is obtained with this set of parameters, where, in average, the rate of symmetric division of type DD is equal to that of the SS type. In other words, in each division, the probability of generating a daughter stem cell is ≃ 0.50. As a result, the number of stem cells in the tissue remains fixed to contribute in future self-renewal and replacement of dead or damaged non-dividing differentiated cells [80].

**Table 1.** The parameters set of the model in Homeostasis

| | $\alpha_{A_{X/Y}}$ | $\alpha_{I_{X/Y}}$ | $\beta$ | $\gamma$ | $n$ | SCB |
| --- | --- | --- | --- | --- | --- | --- |
| two-element switch | 35 | 35 | 45 | 0.38 | 4 | 0.5 |
| four-element switch | 35 | 35 | 47.3 | 0.38 | 4 | 0.5 |
| six-element switch | 35 | 35 | 48.5 | 0.38 | 4 | 0.5 |

The parameters of our ODE model (Equation 1) are set through a grid search. It is assumed that 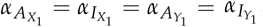. The simulation starts with 1000 populations, each containing of 50 cells, at *t* = *t*_0_, each cell with a two-element regulatory switch, and stops at *t* = *T*. To compute the time evolution of the cell populations through stochastic simulations, we used the Gillespie algorithm[40]. The stem cell birth rate (SCB), the probability of generating stem cells in each cell division, is computed at the end of the simulation and represented in Table 1, the first row, the last column. As seen, with this parameter setting, one-half of the daughter cells remain as stem cells and the other half differentiates and maintains the proliferation/differentiation balance. The simulation is repeated on other populations of cells with four- and six-element internal switches (Equations 11, and 12) and the corresponding parameters are represented in Table 1, the second and third row, respectively.

### B. The perturbation effect in the model

Although a two-element switches could account for describing the interactions between determinants in a cell, it is too sensitive to perturbations which arise from genetic mutations. To be specific, a perturbation in a cell internal switch affects the number of determinants in the cell that could influence the bias of the daughter cells fate toward cell proliferation or differentiation. As a result it affects the the functionality of tissues and organs in replacing damaged or dead cells. A mutation in the system may follows an imbalance in SS and DD division rates which can lead to two different scenarios. First, if the SS division rate surpasses the DD division rate, the stem cells increase in number exponentially. Second, in contrast if the DD division rate surpasses the SS division rate, as time passes, the number of stem cells decreases and there will not be enough number of them to supply the differentiated cells and maintain the functionality of tissues.

To study the model behaviour in the face of perturbations, we examined the effect of mutations in the system. To this purpose, the fluctuations of 1000 populations, each population containing of 50 cells, are simulated by an eleven-phase simulation as it is presented in Figure 6. The mutations occur only in certain number of cells, not in every cell in an adult tissue. Therefor, in the simulations it is assumed that we only study the population of the cells (50 cells in a hypothetical adult tissue) with the chance for mutations in the genes linked to the their internal switch.

**Fig. 6.**
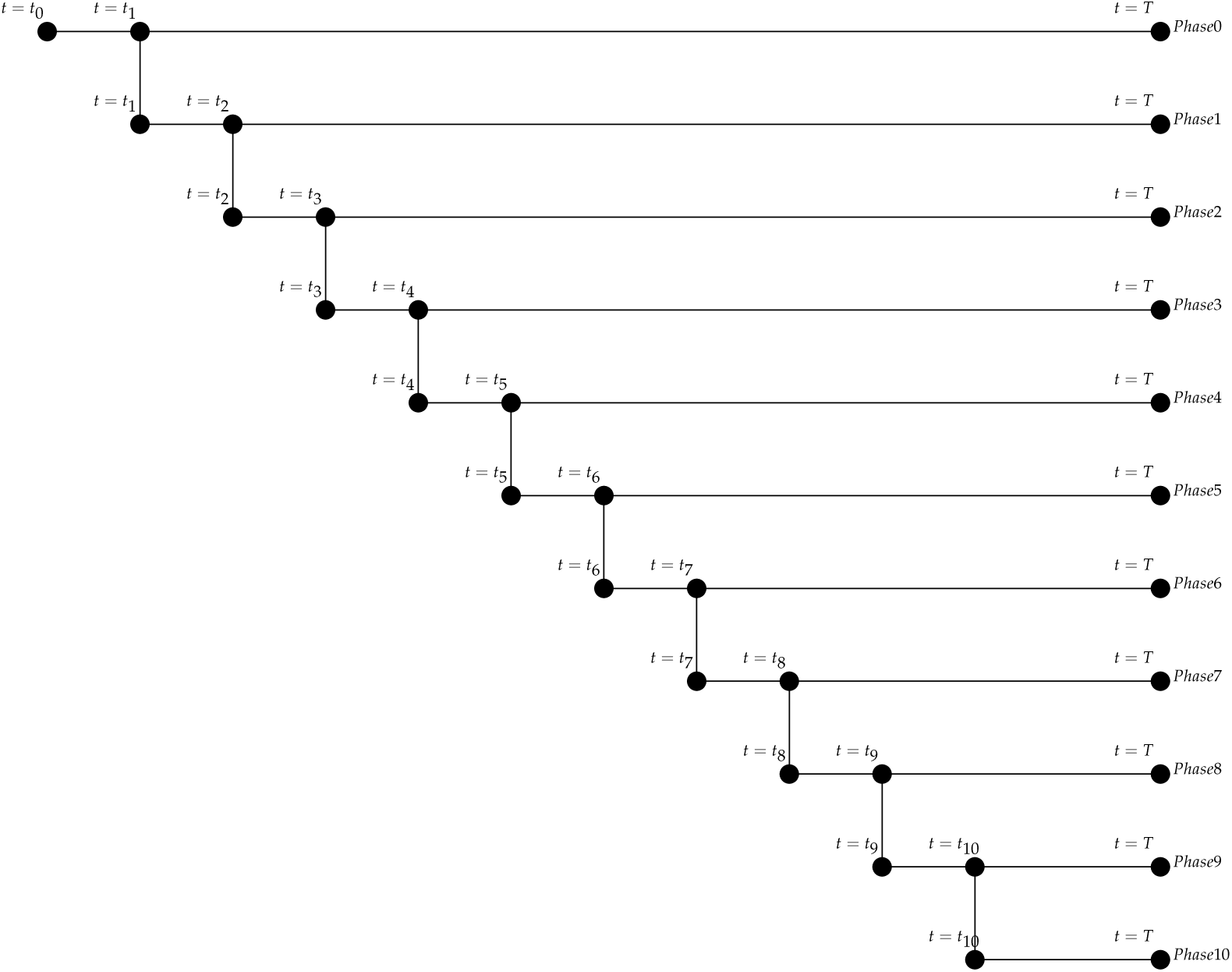
An eleven-phase simulation to study the effect of mutations in the model. Phase 0 starts at *t* = *t*_0_ with 1000 populations, each population containing of 50 cells, and each cell with an internal two-element regulatory switch. Computing the time evolution of the cell populations, stochastic simulations is applied, using Gillespie algorithm. In phase 0, the state of the system in *t* = *t*_1_ when all of the cells have gone through at least two divisions, is stored. Phase 1 starts at *t* = *t*_1_, the time point at which the first mutation occurs to the parameters of the cells’ internal switches. In the same manner, in phase *k* (*k* = 2, 3, …, 10) the state of the system in *t* = *t*_*k*+1_, when all of the cells have gone through at least *k* * 2 divisions, is stored. Phase *k* + 1 starts at *t* = *t*_*k*+1_, the time point at which the *k*^*th*^ mutation with the probability value of *p* occurs to the parameters of the cells’ internal switches. All phases finish at *t* = *T* and the average rate of SS divisions is computed at the end of each phase. Similar simulations are performed for four- and six-element internal switches.

Phase 0 starts at *t* = *t*_0_ and each population cell contains an internal two-element regulatory switch shown in Figure 1.a. The model parameters are chosen from Table 1, the first row. To compute the time evolution of the cell populations, a stochastic simulation, using Gillespie algorithm, is applied. In phase 0, the state of the system in *t* = *t*_1_, when all the cells have gone through at least two divisions, is stored. Then, phase 1 starts at *t* = *t*_1_, the time point at which the first mutation occurs to the parameters of the cells’ internal switches. In the same manner, in phase *k* (*k* = 2, 3, …, 10) the state of the system in *t* = *t*_*k*+1_, when all of the cells have gone through at least *k* * 2 divisions, is stored. Then, phase *k* + 1 starts at *t* = *t*_*k*+1_, the time point at which the *k*^*th*^ mutation occurs to the parameters of the cells’ internal switches.

All eleven phases finish at *t* = *T*. Similar simulations are performed for four- and six-element internal switches (shown in figures 1.b and 1.c) with corresponding parameters set in Table 1, the second and third row.

In our model, mutations are represented with a random change, *ϵ*, in the value of the switch parameters, as following:

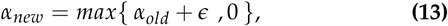

where,

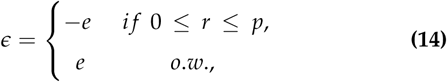

with *r* ∼ *U*(0, 1). We assume that mutations in the cell internal switches rarely change the interaction to be stronger (positive-value mutations), while the majority of them either cause no significant changes in the cell functionality or lead to weaken the interactions (negative-value mutations). Considering these assumptions, in our model, *e* ∼ *Exp*(*λ*) and the probability of negative-value mutations, *p*, is chosen from the set {0.85, 0.90, 0.95, 0.99}. At time *t* = *t*_*k*_ (*k* = 2, 3, …, 10), in each population cell, one of the parameters 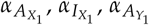, and 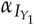 (*α*_*old*_) is randomly chosen and mutated (*α*_*new*_) based on Equation 13. It is also worth mentioning here that as the system behavior in the time interval [*t*_0_, *t*_*k*_] is similar in phase 1 and *k* (*k* = 2, 3, …, 10), without loss of generality, one can say that all phases start from *t* = *t*_0_.

At the end of each phase, we computed the number of populations resisting the perturbations (Figure 7), as well as the probability of generating a stem daughter cell per cell division (Figure 8). Both figures contain four subplots corresponding to the probability values of *p* = 0.85, *p* = 0.90, *p* = 0.95, and *p* = 0.99. Each subplot demonstrates three curves for two-, four-, and six-element switches. The results show that by increasing the number of mutations, the number of populations which last to the end of the simulation decreases, while the stem cell birth rate (SCB) increases.

**Fig. 7.**
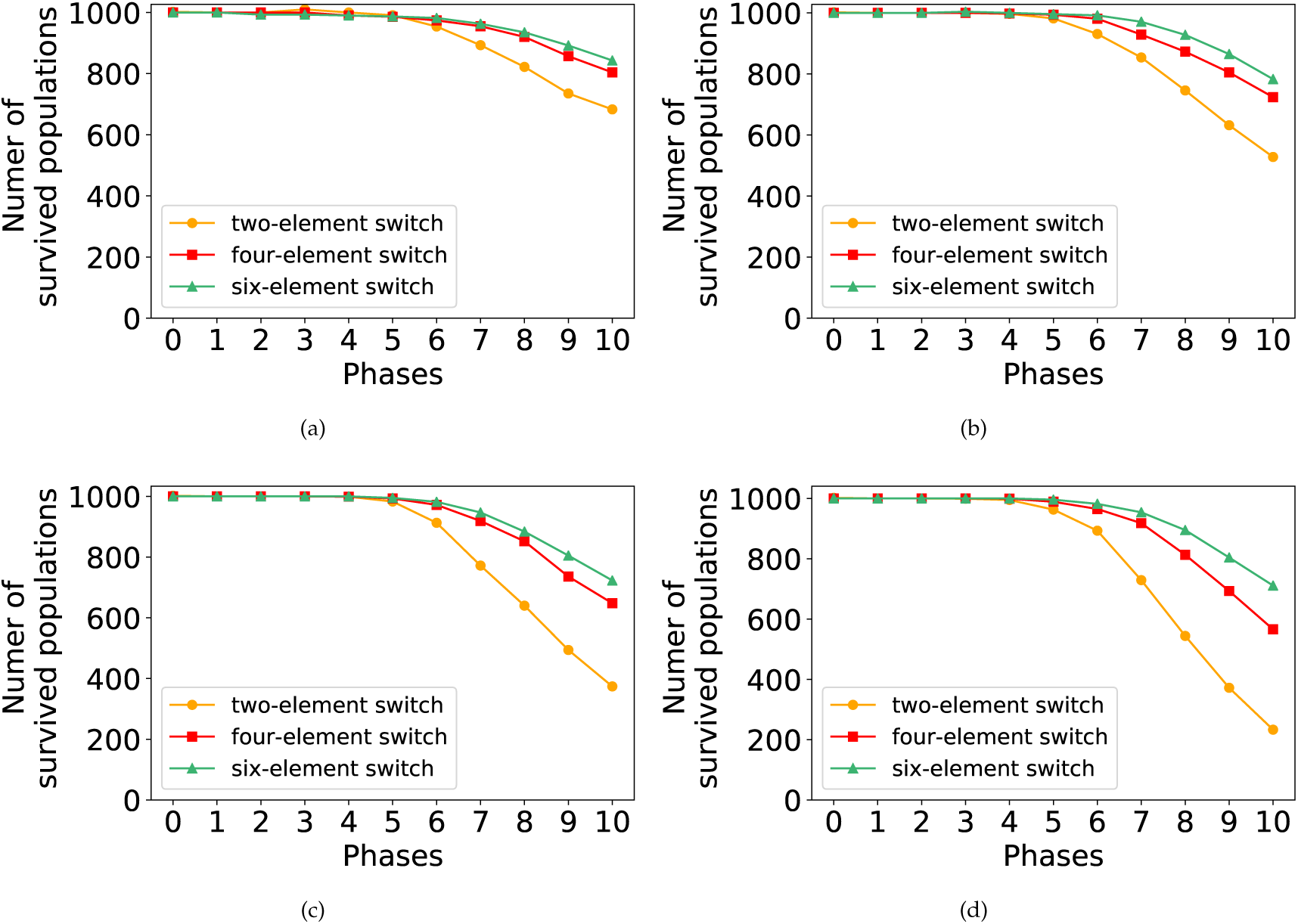
Number of populations survived in the face of mutations through the eleven phases of the simulation with the probability value of *p* = 0.85, *p* = 0.90, *p* = 0.95, and *p* = 0.99 corresponding to subplots (a), (b), (c), and (d), respectively.

**Fig. 8.**
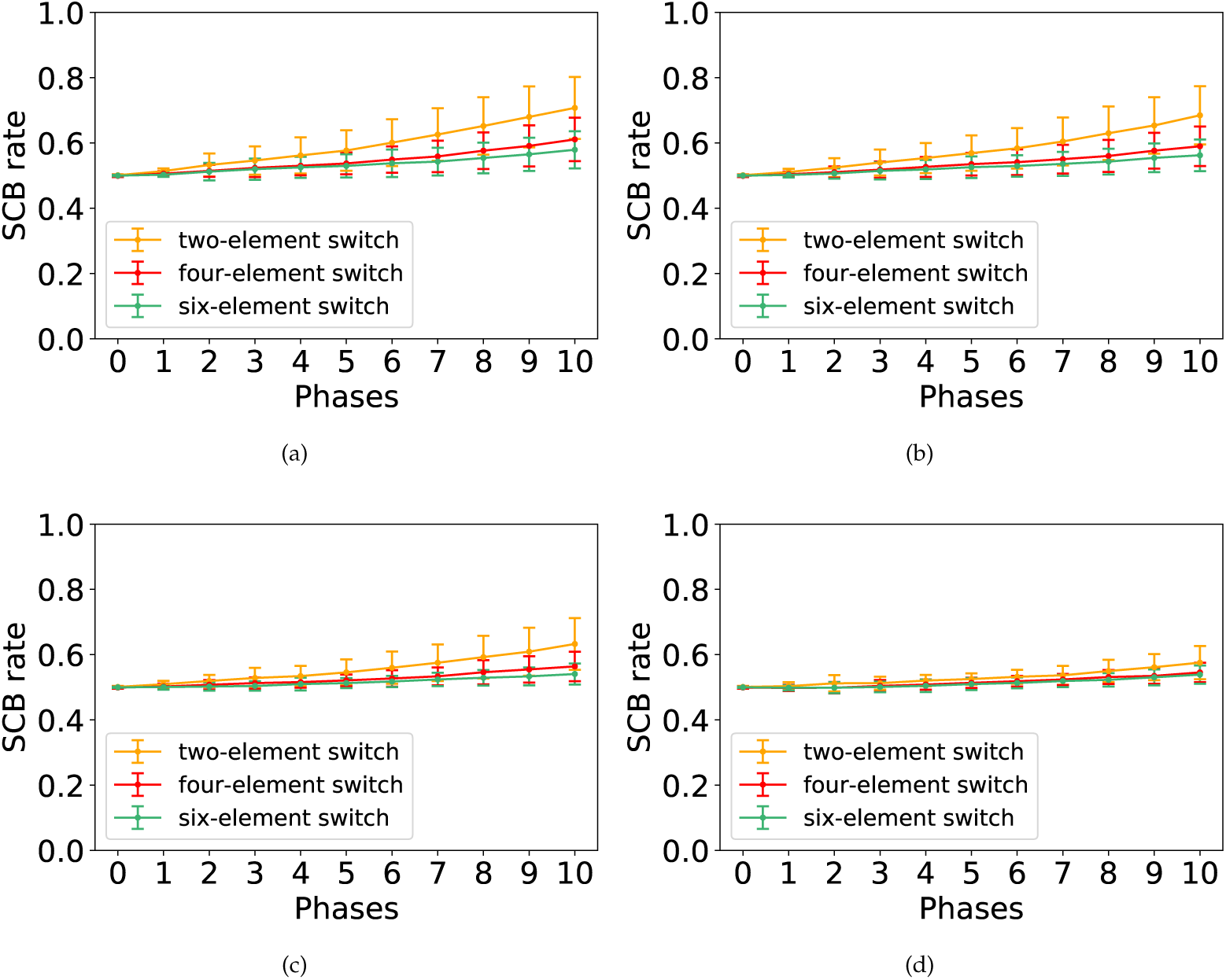
SCB rate in the face of mutations through the eleven phases of the simulation with the probability value of *p* = 0.85, *p* = 0.90, *p* = 0.95, and *p* = 0.99 corresponding to subplots (a), (b), (c), and (d), respectively.

The bias of the cell internal switches is determined by the mean and variance of determinants around the middle attractor. The parameters in Table 1 determine a distribution of determinants which guarantees the proliferation/differentiation balance. By the defined distribution, after each cell division, half of the daughter cells remain in yellow region of Figure 4 (as stem cells), while the other half are born in the green or orange regions of Figure 4 (as differentiated cells).

Any mutation in the model parameters increases the variance around the middle attractor. As a result, two scenarios are possible. First, in some of the populations most of the stem cells are located close to the boundaries of the three-region force-field representation of Figure 4. Therefore, in the next generation, their daughter cells are more probable to be born in the green or orange regions (as differentiated cells). In the other word, it shifts the differentiation/proliferation balance toward differentiation and increases the population extinction probability. This explains why the number of populations declines in the face of mutations (Figure 7). In the second scenario, in some of the populations most of the stem cells are placed far from the boundaries and the original middle attractor. Therefore, in the next generation, their daughter cells are more likely to remain in the yellow region of Figure 4 (as stem cells). In contrast with the first scenario, it shifts the differentiation/proliferation balance toward proliferation and increases the population growth rate (SCB rate). This is compatible with the system behavior in the face of mutations shown in Figure 8.

In each subplot of Figures 7 and 8, it is obvious that four- and six-element switches could better buffer the perturbations effects compared to the two-element switches. It is in agreement with the great idea of Waddington, “canalisation” in [79]. Besides, the curves in Figure 7 show that the perturbation effect on the number of populations surviving to the end of simulations is more pronounced when the value of *p* is relatively large. In contrast, the model perturbation effect on SCB rate is weaker for the large values of *p* (Figure 8).

To analyze the effect of mutations on the number of populations last to the end of phase 10 (Figure 7) and on the SCB rate (8), it is necessary to study the dynamics of our model through Figures 9, S1, S2, S3, S4, S5, S6, S7, S8, S9, S10, S11.

**Fig. 9.**
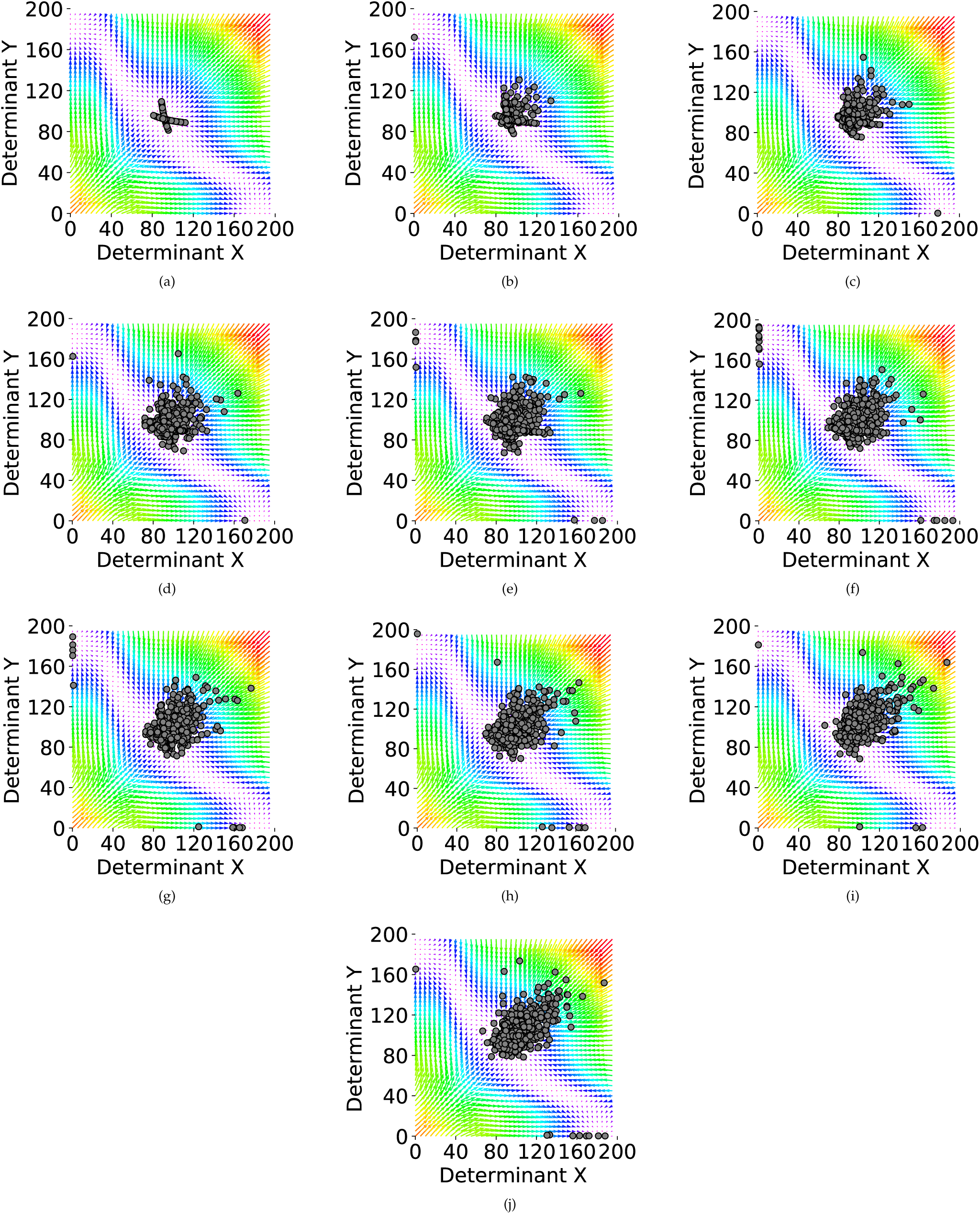
Ten phases of the simulation with the probability value of *p* = 0.85, and *λ* = 5. The internal regulatory networks of cells are assumed to be two-element switches. (a-j) Phases 1 to 10 of the simulations. In each one of the plots, each circle represents the middle attractor of one of the cells in the population, with the representative cell being the one which produces the highest proportion of stem daughter cells at the end of each phase.

The force-field representations of our tristable model corresponding to each of the ten simulation phases (phase 1, 2, …, 10) with two-, four-, and six-element switches are demonstrated in Figures 9, S1, and S2, respectively. For these figures, it is assumed that *p* = 0.85 and *λ* = 5.

In each subplot, each circle represents the middle attractor of one of the cells in the population. The representative cell is the one which produces the highest proportion of stem daughter cells at the end of each phase. It is clear that the number of circles in each subplot is equal to the number of populations last to the end of each phase. In the same manner, Figures S3, S4, and S5, Figures S6, S7, and S8, and Figures S9, S10, and S11, respectively demonstrate the dynamics of our model for the probability value of *p* = 0.90, *p* = 0.95, *p* = 0.99.

By a single mutation with a positive (negative) value of *ϵ* (Equation 14), the corresponding middle attractor tends toward the upper(lower) triangular portion of force-field representation (Figures 9.a, S1.a, S2.a, S3.a, S4.a, S5.a, S6.a, S7.a, S8.a, S9.a, S10.a, S11.a). As mentioned previously, the growth rate of a single cell increases when its corresponding middle attractor is far from the boundaries and the original middle attractor. Since for the larger values of *p* (*p* = 0.95, and *p* = 0.99), positive-value mutations are less probable, most of the mutated cells attractors are located close to the boundaries and the original middle attractor (Figures S6, S7, S8, S9, S10, S11). Therefore, most of the populations vanish in the face of mutations (Figure 7). In contrast, for the smaller values of *p* (*p* = 0.85, and *p* = 0.90), positive-value mutations are more probable, and most of the mutated cells attractors are located far from the boundaries and the original middle attractor in the direction of the minor diagonal of the force-field representation (Figures 9, S1, S2, S3, S4, S5). Therefore, the growth rate is easily affected by mutations for the smaller values of *p* (Figure 8).

Studying the dynamics of our model through ten subplots of each of the Figures 9, S1, S2, S3, S4, S5, S6, S7, S8, S9, S10, S11 reveal how populations facing a single mutation behave differently from the ones facing the accumulation of mutations. For the ease in discussion, since there is a one-to-one correspondence between the cells and their middle attractor, we assume that each circle represent the cell which produces the highest proportion of stem daughter cells in the population.

When a single mutation occurs in a cell, either the cell would be located close to the boundaries and dies through the next division or it would be located far from the boundaries with a higher growth rate and remains in the population. In the same manner, in the next generation, when a single mutation occurs in a survived cell, either the cell would be located close to the boundaries and dies through the next division or it would be located far from the boundaries with a higher growth rate and remains in the population^1^. This process is repeated for all future mutations. One can say, if a cell remains in the population and receives the 10^*th*^ mutation, it is a cell with a high growth rate (with a great chance), i.e. a cell which is far from the boundaries in the direction of the minor diagonal of the force-field representation.

Each subplot in Figure 10 shows the rate of stem cell birth in 50 cells with the highest growth rate, 50 cells among all the populations cells resisting the perturbations to the end of each phase. Figure 10 illustrates that accumulation of mutations could give rise to the birth of cells which always divides symmetrically to produce two daughter stem cells, cells with the SCB rate value of ≃ 100%. In other words, mutation accumulation results in the birth of “immortal cells” which pass through several symmetric divions which lead to exponemtially growth in population number (see Figure 11). Each row in Figure 11 shows the population size distribution among eleven phases of the simulation for two-, four-, and six-element switches, respectively.

**Fig. 10.**
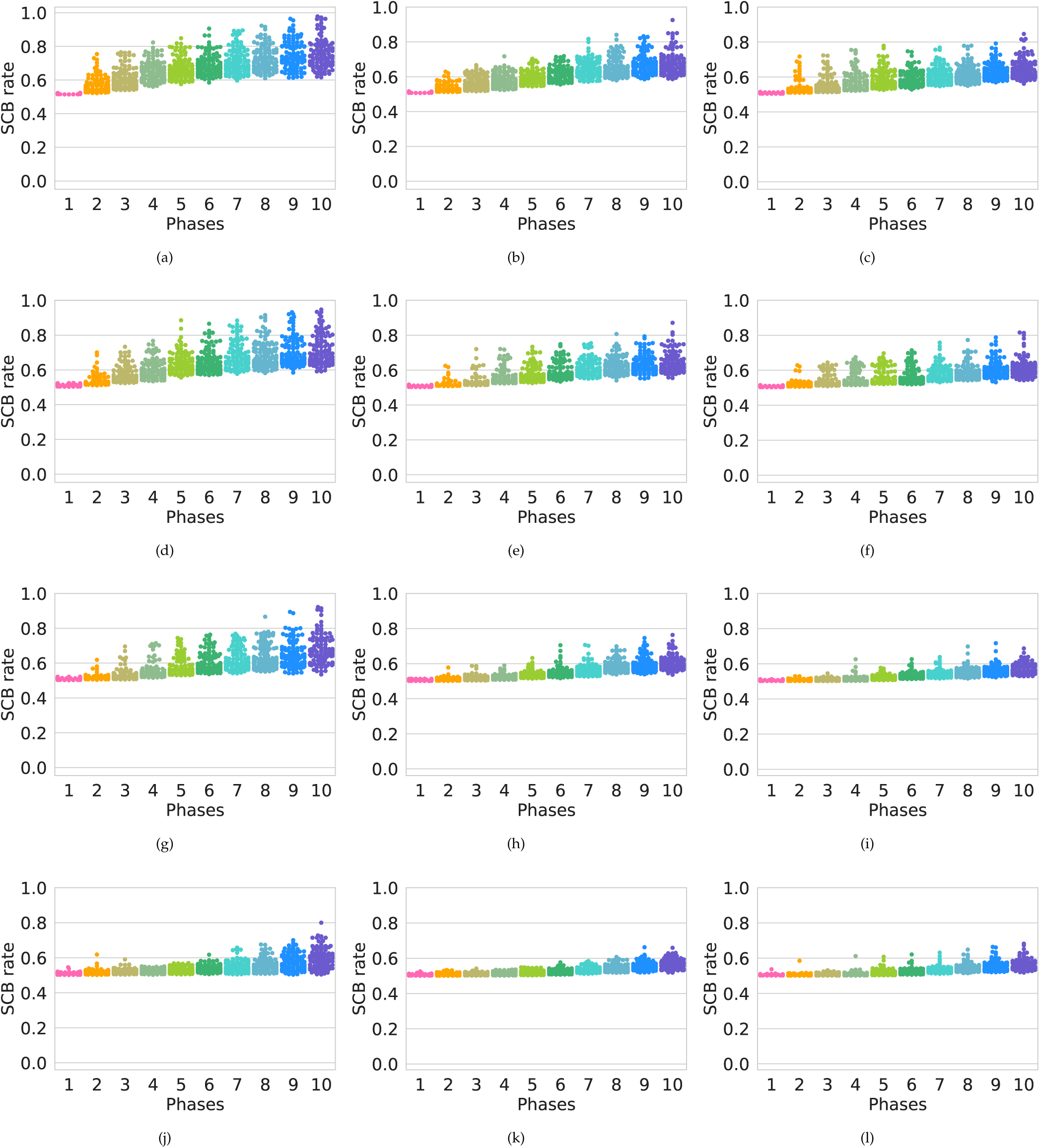
Swarm plot of the SCB rate in ten phases of the simulation. (a, b, c) Swarm plot for the populations of cells with two-element, four-element, and six-element switches, respectively with the probability value of *p* = 0.85. (d, e, f) Swarm plot for the populations of cells with two-element, four-element, and six-element switches, respectively and the probability value of *p* = 0.90. (g, h, i) Swarm plot for the populations of cells with two-element, four-element, and six-element switches, respectively and the probability value of *p* = 0.95. (j, k, l) Swarm plot for the populations of cells with two-element, four-element, and six-element switches, respectively and the probability value of *p* = 0.99.

**Fig. 11.**
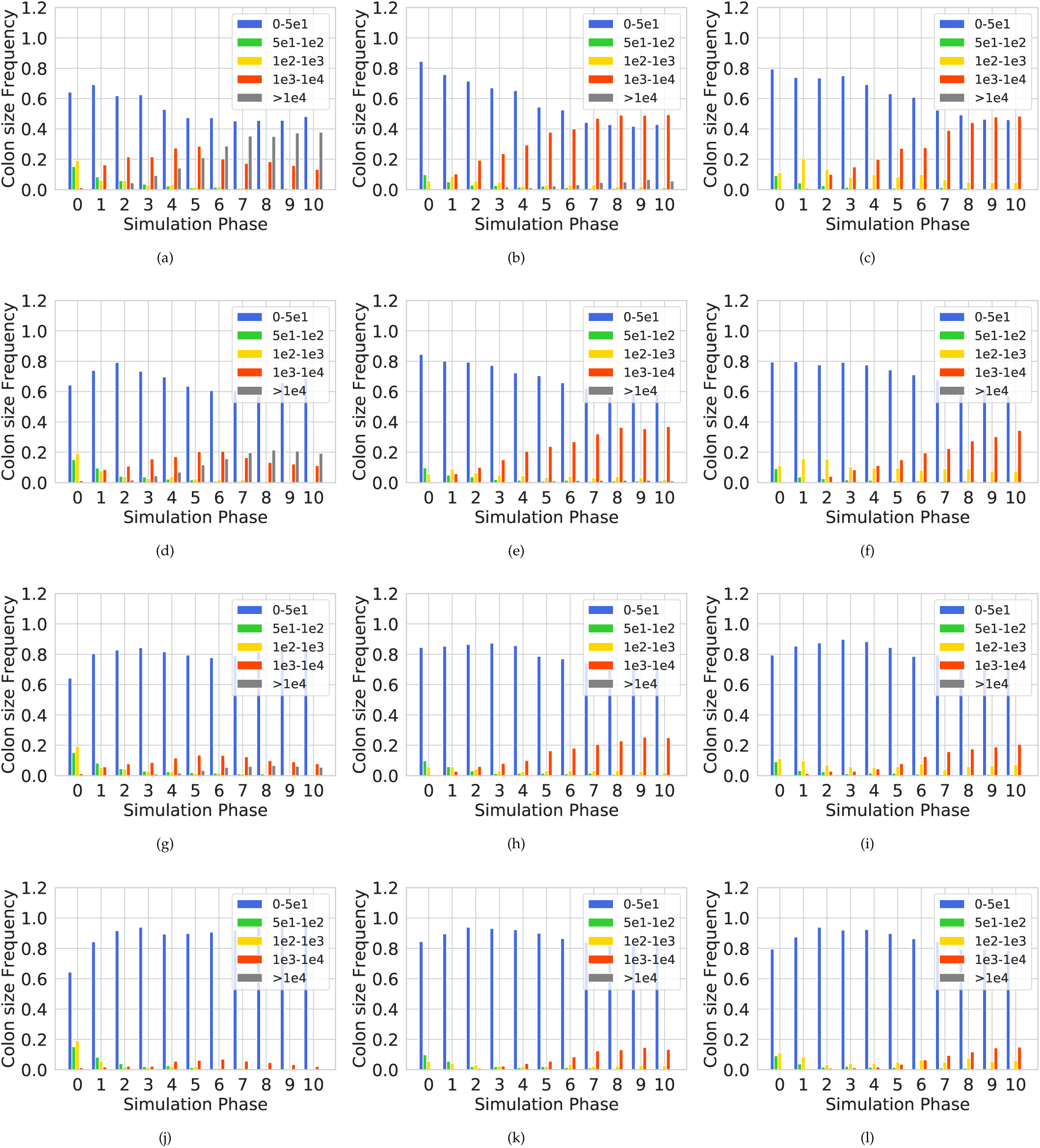
Population size distribution among eleven phases of the simulation. (a,b,c) The population of cells with two-element, four-element, and six-element switches, respectively and the probability value of 0.85%. (c,d,f) The population of cells with two-element, four-element, six-element switches, respectively and the probability value of 0.90%. (g,h,i) The population of cells with two-element, four-element, six-element switches, respectively and the probability value of 0.95%. (g,h,i) The population of cells with two-element, four-element, six-element switches, respectively and the probability value of 0.99%.

In both Figures 10 and 11, it is clearly seen again that more complex switches provide more robustness, and that for the larger values of *p*, cells with the high rate of SCB are less probable while the great number of cell populations undergo a decline contrasting with the smaller values of *p*.

We have designed two eleven-phase simulations corresponding to two different values, *λ* = 2, and *λ* = 10 (Equations 13, and 14). Similar to the simulation which was described previously (Figure 6), simulations start at *t* = *t*_0_ and stop at *t* = *T*, with 1000 populations with 50 cells, where each cell contains a two-element internal switch, and *p* = 0.95. Figure 12 shows how our model behaviour is influenced by the values of the parameter *λ* (*λ* = 2, *λ* = 5, and *λ* = 10). The dynamics of our system through ten phases of the simulations is shown in Figures S12, and S13 for *λ* = 2, and *λ* = 10, respectively. Analyzing Figures 12, S12, and S13 reveals that perturbations with *λ* = 2 merely can affect the system behaviour, while perturbations with *λ* = 10 exhibit high random variation in the system behaviour.

**Fig. 12.**
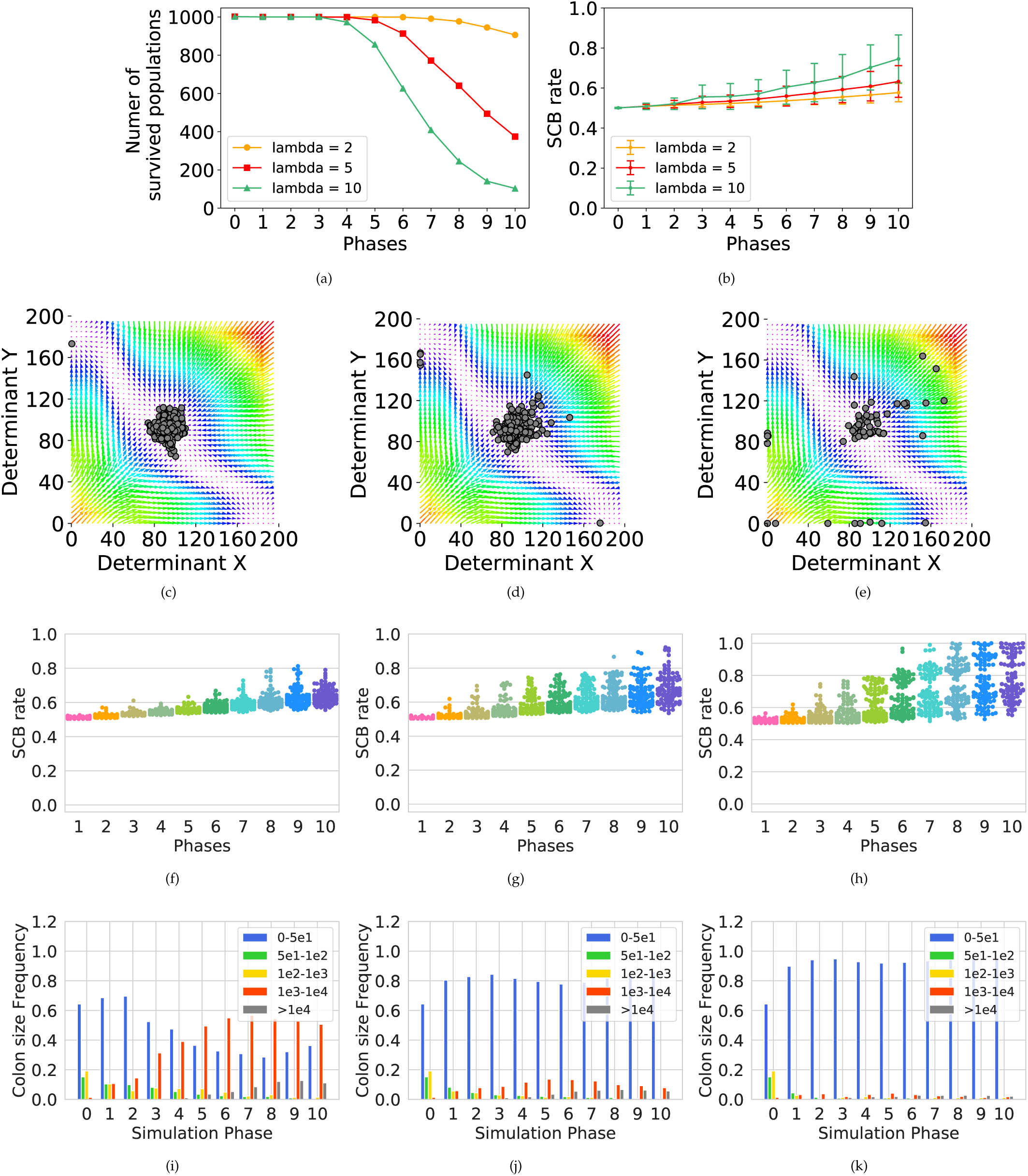
The system behaviour in the face of mutations through the eleven phases of the simulation with different values of parameter. *λ*. The internal regulatory networks of the cell populations are two-element switches, and the probability value of *p* is equal to 0.95. (a, b) SCB rate and number of populations survived in the face of mutations. (c, d, e) Phases 10 of the simulations with *λ* = 2, *λ* = 5, and *λ* = 10, respectively. In each one of the plots, each circle represents the middle attractor of one of the cells in the population, with the representative cell being the one which produces the highest proportion of stem daughter cells at the end of each phase (f, g, h) Swarm plot of the SCB rate with *λ* = 2, *λ* = 5, and *λ* = 10, respectively. (i, j, k) Population size distribution among eleven phases of the simulation with *λ* = 2, *λ* = 5, and *λ* = 10, respectively.

### C. Changing the bias of switch by changing the parameters

The proportion of cells which remain as stem cells to continue self-renewal or that which begin the the pathway to differentiation is clearly related to the area of three attractors domain (Figure 4). In other words, the final fate of the daughter cells could be influenced by the value of the parameters in equation 1. Besides, depending on the intensity of inhibitory and activatory effects of determinants (through the values of constants in the Hill function ([27]), the attractor domains could be symmetric or not. Figure 13 shows the system behaviour in parameter space by evaluation of the effect of six parameters (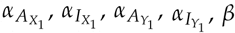, and *γ*) of equation 1 in six columns. The middle row indicates *X* − *Y* phase plane corresponding to the original set of parameters of Table 1, the first row.

**Fig. 13.**
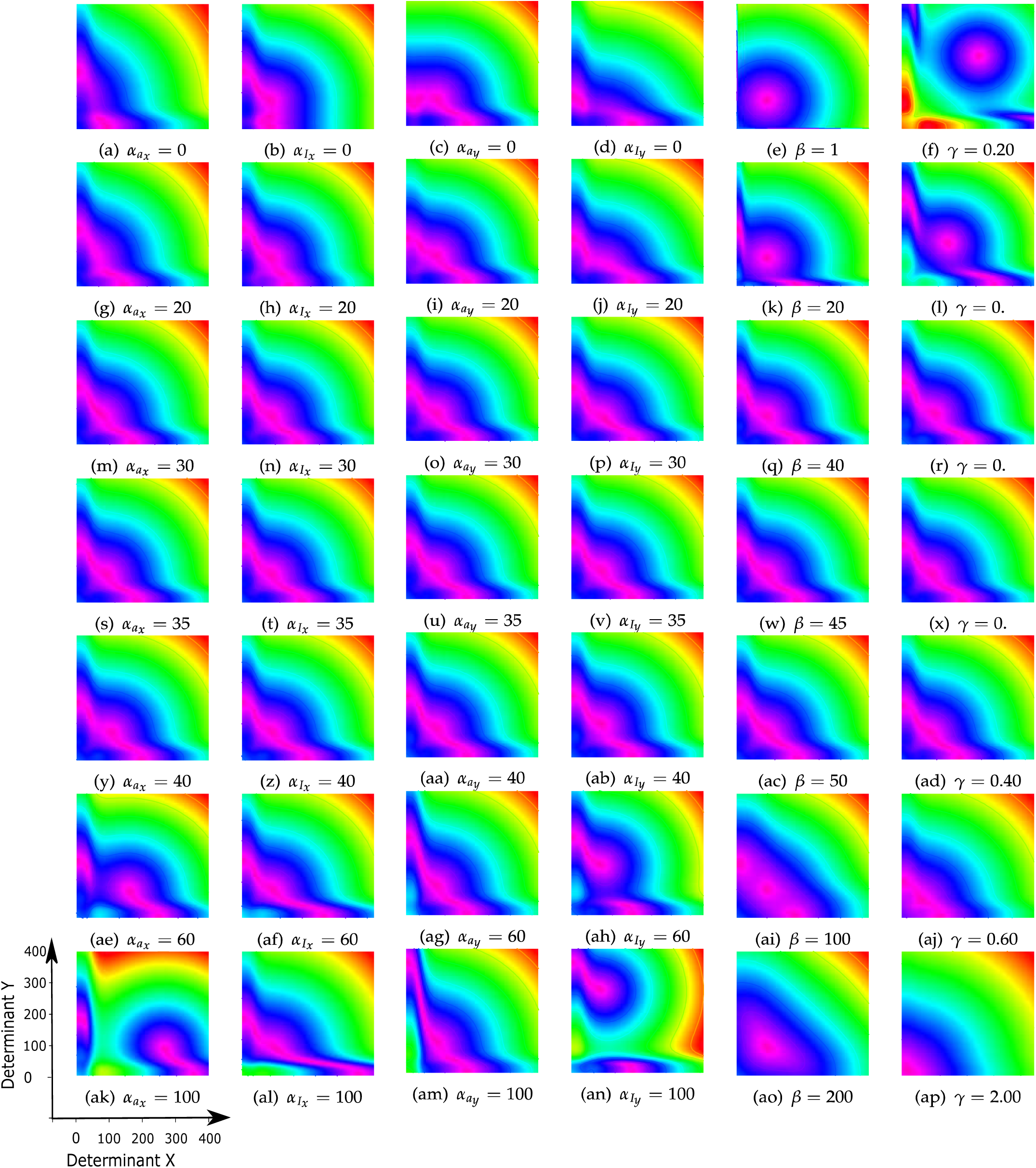
The system behavior in parameter space. Evaluation of the effect of parameters’ changes for equation 1. In the 9 × 6 array, each cell represents the *X* − *Y* phase plane, for parameter values as indicated at the bottom of each subplot. The magenta regions are highly stable. The basic set of parameters are chosen from Table 1, 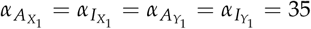, *β* = 45, *γ* = 0.38, and *n* = 4.

Although our model describes a symmetric dynamical system, Figure 13 shows that by changing the parameters in the model it can be used to study any desired tissue with different proportion of differentiated cells. It reflects the flexibility of our model.

## 4. DISCUSSION

By focusing on the effect of stochasticity on the cell final fate, we computationally modeled a regulatory mechanism to control the proliferation/differentiation balance maintaining the homeostatic state in a hypothetical adult tissue. In the most simple model, it is assumed that this hypothetical developing tissue consists of stem cells and two differentiated cell types. Our model has been described by a set of ordinary differential equations to model a regulatory switch (Equation 1). This switch consists of two cytoplasmic cell fate determinants with auto-activation and mutual inhibition (Figure 1.a) which forms a tristable dynamical system. The results showed that two-element switches can be significantly affected by the system perturbations, while the more complex switches (Figures 1.b and 1.c) provide more robustness. This is somehow similar to the idea of “canalisation” in Waddington’s book [79]. Several biological observations being collected to support the existence of internal switches consisting two groups of determinants, with feedback activation within each group and feedback inhibition between the groups [36]. Each dividing stem cell contains a small number of determinants, and a small change could significantly affect the tissue’s final fate. The extended regulatory networks work as a crucial defence against the perturbations in the system. Moreover, our analysis reflects the flexibility of our model to describe any desired tissue with different proportion of differentiated cells (Figure 13).

By defining the noise as the distance between the number of determinants in each cell and the expected number of them in the population (the original middle attractor), the cell determinants and the noise are from the same family of distributions. Therefore, one can say, the cell noise distribution controls the proliferation/differentiation balance in the population to maintain homeostatic state. When the noise variation increases, the majority of the daughter cells are born as differentiated cells (in orange or green region of Figure 4). In this case, after several generations, there wont be enough stem cells to replace the dead differentiated cells in the tissue. This can be interpreted as aging [81]. On the other hand, by decreasing the noise variation, a great number of daughter cells are born as stem cells. Under this condition, the growth rate of the cell population increases through the future generations. This can be interpreted as cancer [82]. This indicates the key role the noise plays in cell decision-making by regulating the probability of differentiation in a normal adult tissue.

To keep a pool of *N* stem cells in an adult tissue, the original stem cells must produce *N* stem cells. For this purpose, each division produces one stem cell and one differentiated cell, on average [3]. Without loss of generality, we can say that in homeostatic state, all the cells produce exactly one stem cell and one differentiated cell (SCB rate in phase 0 from Figures 8). When a mutation occurs in a cell, two following scenarios are possible, stem cell extinctions (Figures 7, and 11) or exponential expansion of them (Figures 8, 10, and 11).

In the former case, the mutated cell produces two differentiated cells through the next division (with a great chance). As a result, our hypothetical tissue contains *N* − 1 stem cells. Two new born daughter cells carry their mother cell’s mutation. However, differentiated cells are non-dividing cells and the inherited mutation will be omitted from the population by their death. Therefore, this mutation does not affect the next generation of the cell population. In other words, the only impact of the mutation on the population is the extinction of one stem cell within the stem cell pool.

In the latter case, the mutated cell divides into two daughter stem cells (with a great chance) which leads to *N* + 1 stem cells in our hypothetical tissue. Two new born daughter cells carry their mother cell’s mutation. In contrast with the former case, stem cells are dividing cells and the inherited mutation not only remains in but also spreads throughout the population via the symmetric cell divisions. Consequently, this mutation results in the exponential expansion of the stem cell pool. It resembles the behaviour of the dividing tumour cells with a strong bias toward generating dividing over non-dividing daughter cells through cell division [83].

In this study, aging is defined as a process through which the tissue gradually loses the stem cells with their self-renewal and regenerative potential. Also, cancer is defined as a process in which an individual mutant pool of stem cells divides and increase in mass, out of control. Based on these definitions and considering the consequences of single and accumulative mutations in the population, it is easily concluded that aging is a slow process while cancer can grow so fast.

Maintaining tissue homeostasis is strongly linked to the stem cell divisions with the risk of mutations in the next generation. In other words, in a long period of time and in a large enough population, mutations are inevitable [84]. When a single mutation occurs, in the genes linked to the cell internal switch, it influences the bias of the daughter cells’ fate toward either cell differentiation (and death) or cell proliferation with a higher growth rate (*g*_1_ > 0.50). In the former case, the mutation will be removed from the population while in the latter case the mutation remains in. In the same manner, in the next generation, when a single mutation occurs in a survived cell, either the cell dies through the next division or remains in the population with a higher growth rate (*g*_2_ ≥ *g*_1_ > 0.50). This process is repeated for all future mutations. One can say, if a cell remains in the population and receives more mutation, it is a cell with a high growth rate (with a great chance). These dividing cells, with the accumulation of inherited mutations, undergo symmetric cell divisions which lead to exponential expansion of the stem cell pool in the tissue (Figures 8, and 11). It reveals how populations facing a single mutation behave differently from the ones facing the accumulation of mutations.

As it is mentioned, mutation accumulation can result in developing cancer. It clearly explains how cancer can be considered as an age-related process. In the other words, as a sufficiently long period of time is needed to grow mutated cells which multiply in great number, one can say the probability of cancer incidence increases with age [84, 85]. In addition, this is in keeping with those studies emphasizing on the importance of the total number of the stem cell divisions, to receive successive mutations, in the lifetime risk of many cancer types [84–86]. Moreover, it can be easily concluded that if someone is born with inherited genetic mutation, it puts them at a higher risk of cancer.

In tissue homeostasis, SCB rate after each stem cell division is equal to 0.50 on average. On the other words, all stem cells have similar capacity to self-renew and/or differentiate. However, analysing Figure 11 (in phase 0) reflects the fact that stem cells behave in a stochastic manner when they are studied individually. Although the size of the stem cell pool remains fixed in the tissue (Phase 0 from Figure 8), some populations shrink whereas some others expand in size [5]. Figure 11 shows that the population behaviour can be described as a gambling game with equal odds as it was discussed in [19]: *‘equal chance’ does not guarantee ‘equal outcome’*. It implies how the presence of controlled noise in a population of genetically similar cells with the same environmental condition provides both heterogeneity and homeostasis [6].

## ACKNOWLEDGMENTS

The authors thank Steffen Rulands for insightful discussions, and Saeed Reza Kheradpishe, and Aboutaleb Amiri for their valuable suggestions in the preparation of this paper. The first author was partly supported by a grant from IPM. For this project, we made use of high-performance computing clusters, provided by the MPI-PKS.

## Footnotes

1 Clearly, it is possible that for a specific cell, the growth rate is not increased by occurring the second mutation. However, in this case the cell dies in the next generation with a great chance. Therefore, without loss of generality we assume that we are studying the cells with increasing growth rate.

