## Supplementary material for "A Computational Model of Stem Cell Molecular Mechanism to Maintain Tissue Homeostasis": Supplimental File

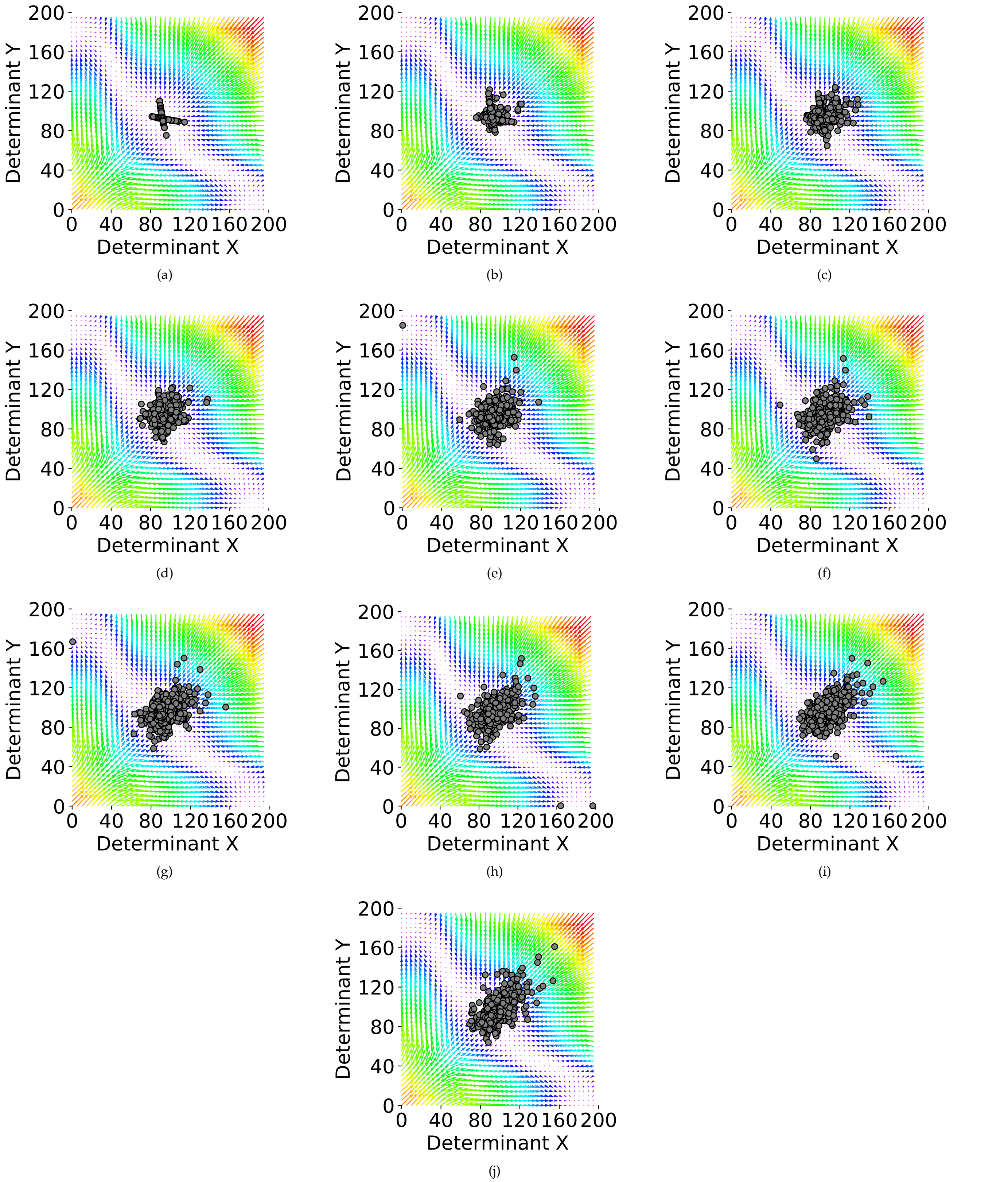


**Fig. S1. Ten phases of the simulation with the probability value of $p = 0.85$, and $\lambda = 5$. The internal regulatory networks of cells are assumed to be four-element switches.** (a-j) Phases 1 to 10 of the simulations. In each one of the plots, each circle represents the middle attractor of one of the cells in the population, with the representative cell being the one which produces the highest proportion of stem daughter cells at the end of each phase.

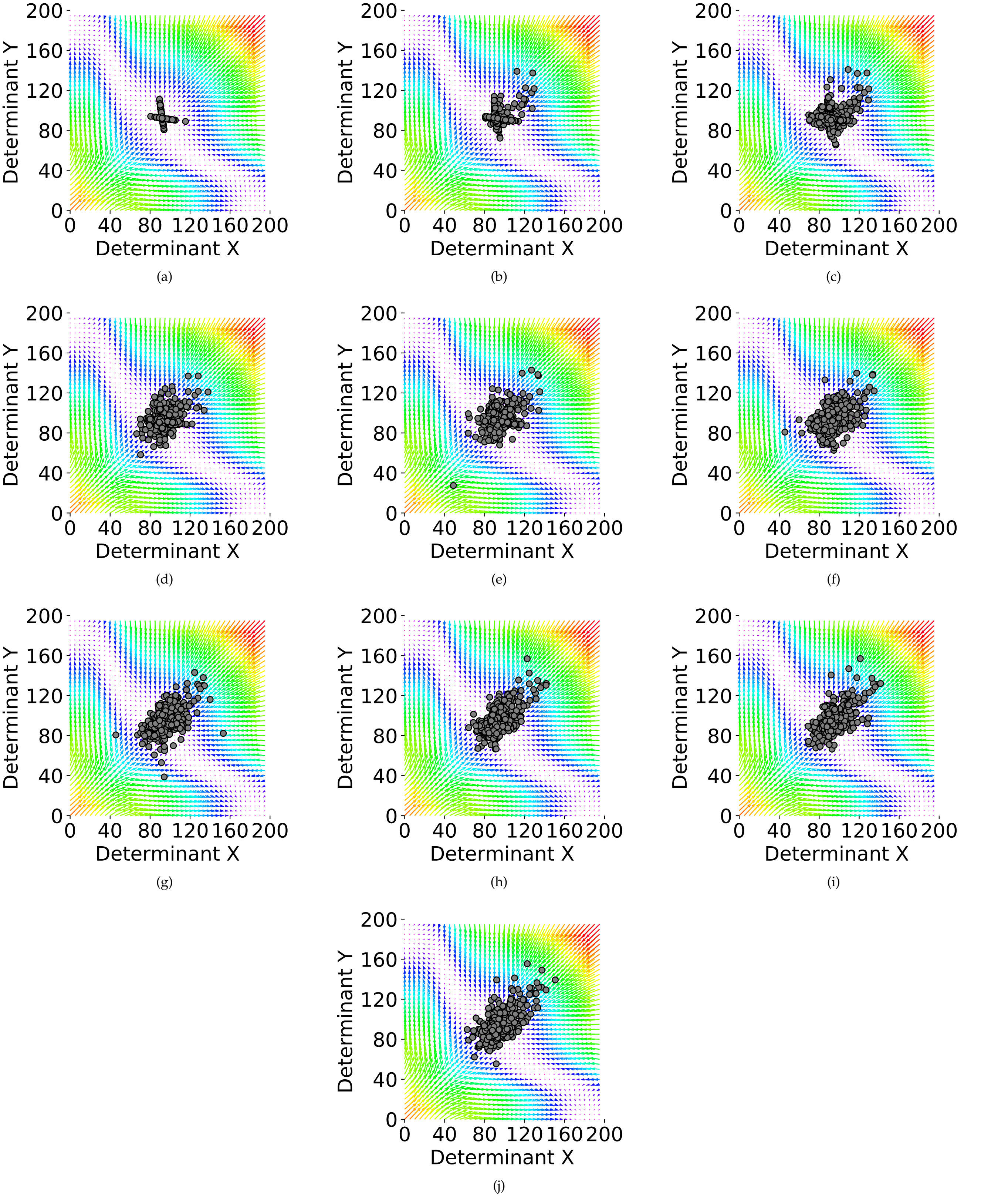


**Fig. S2. Ten phases of the simulation with the probability value of $p = 0.85$, and $\lambda = 5$. The internal regulatory networks of cells are assumed to be six-element switches.** (a-j) Phases 1 to 10 of the simulations. In each one of the plots, each circle represents the middle attractor of one of the cells in the population, with the representative cell being the one which produces the highest proportion of stem daughter cells at the end of each phase.

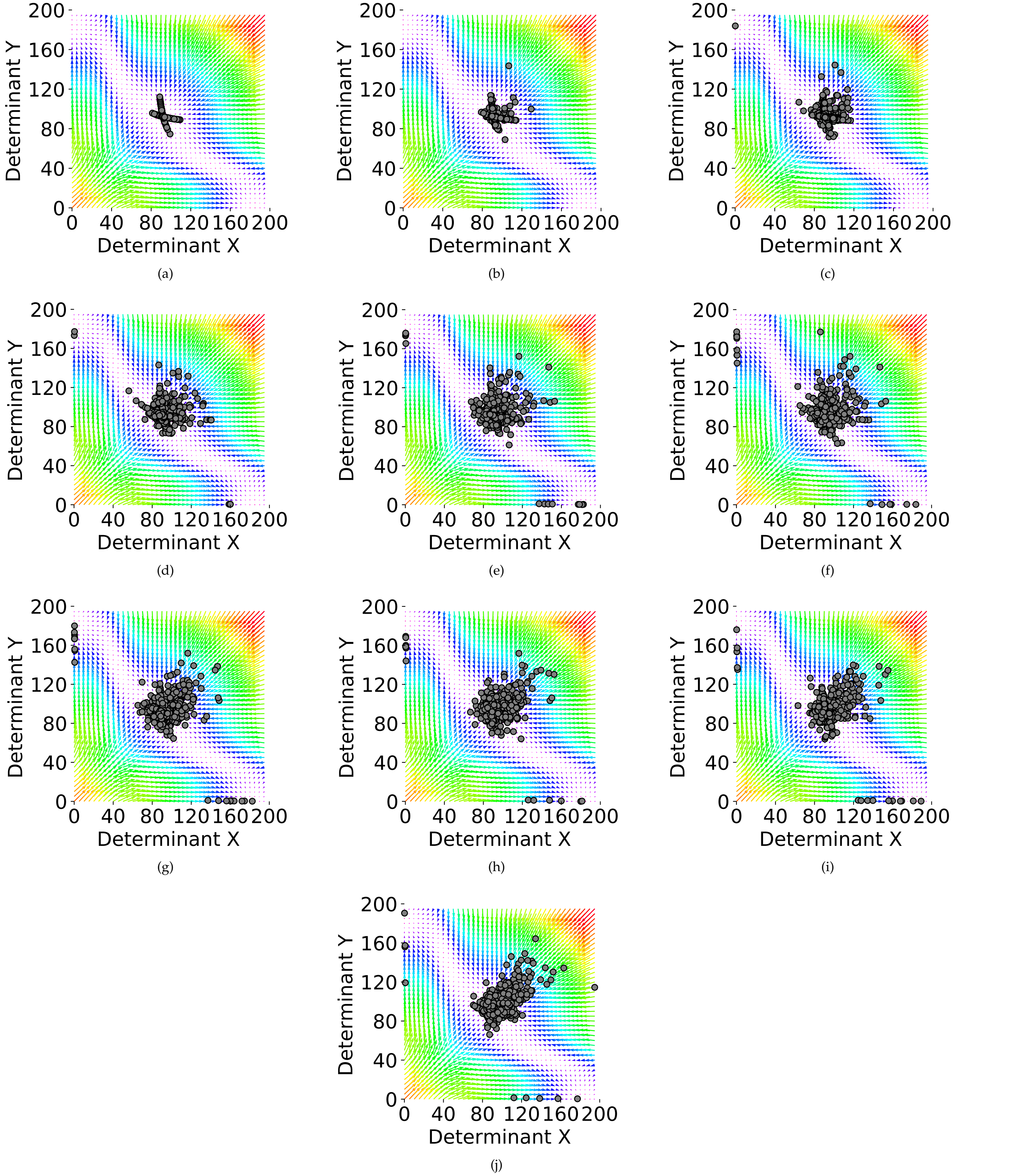


**Fig. S3. Ten phases of the simulation with the probability value of $p = 0.90$, and $\lambda = 5$. The internal regulatory networks of cells are assumed to be two-element switches.** (a-j) Phases 1 to 10 of the simulations. In each one of the plots, each circle represents the middle attractor of one of the cells in the population, with the representative cell being the one which produces the highest proportion of stem daughter cells at the end of each phase.

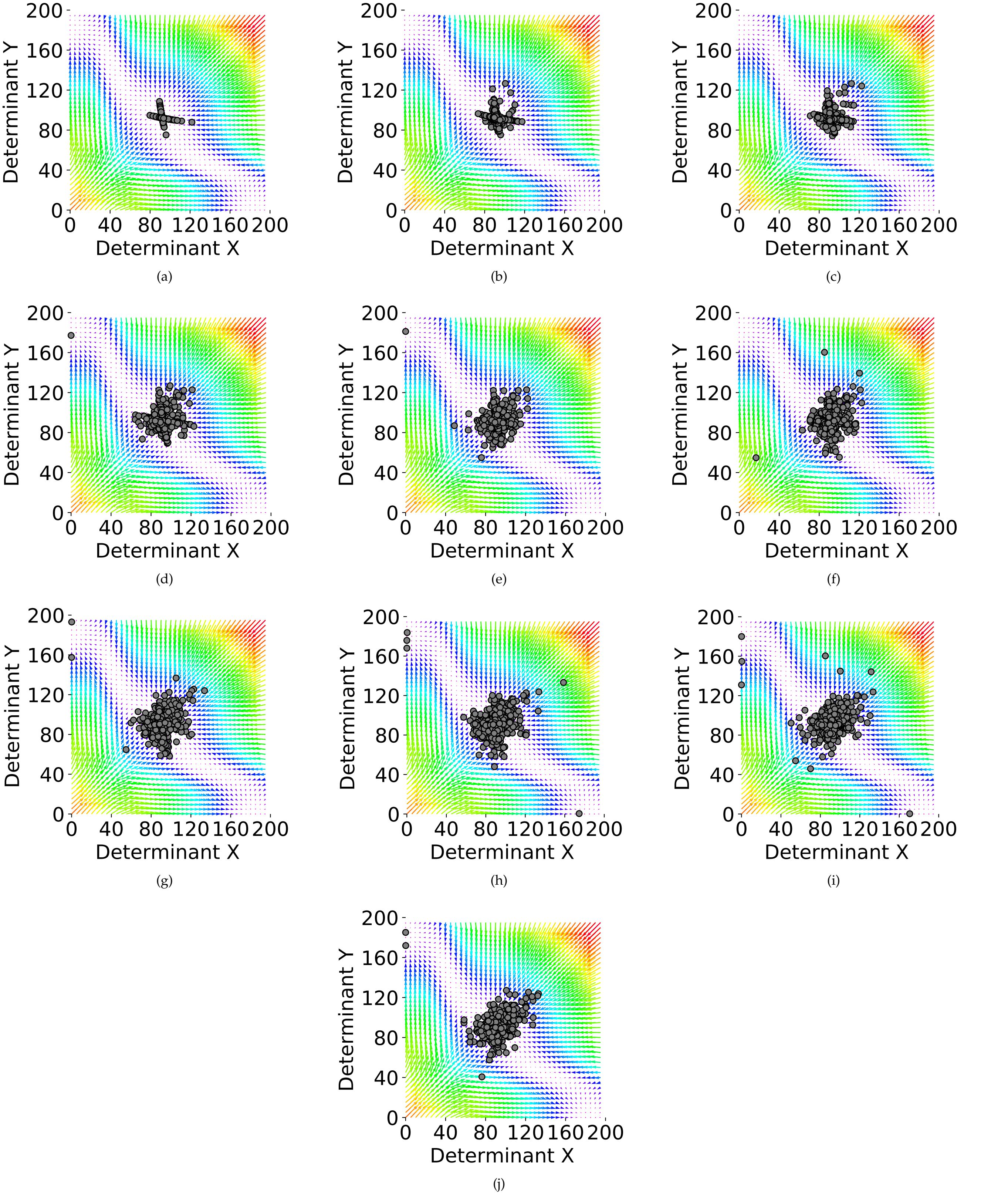


**Fig. S4. Ten phases of the simulation with the probability value of $p = 0.90$, and $\lambda = 5$. The internal regulatory networks of cells are assumed to be four-element switches.** (a-j) Phases 1 to 10 of the simulations. In each one of the plots, each circle represents the middle attractor of one of the cells in the population, with the representative cell being the one which produces the highest proportion of stem daughter cells at the end of each phase.

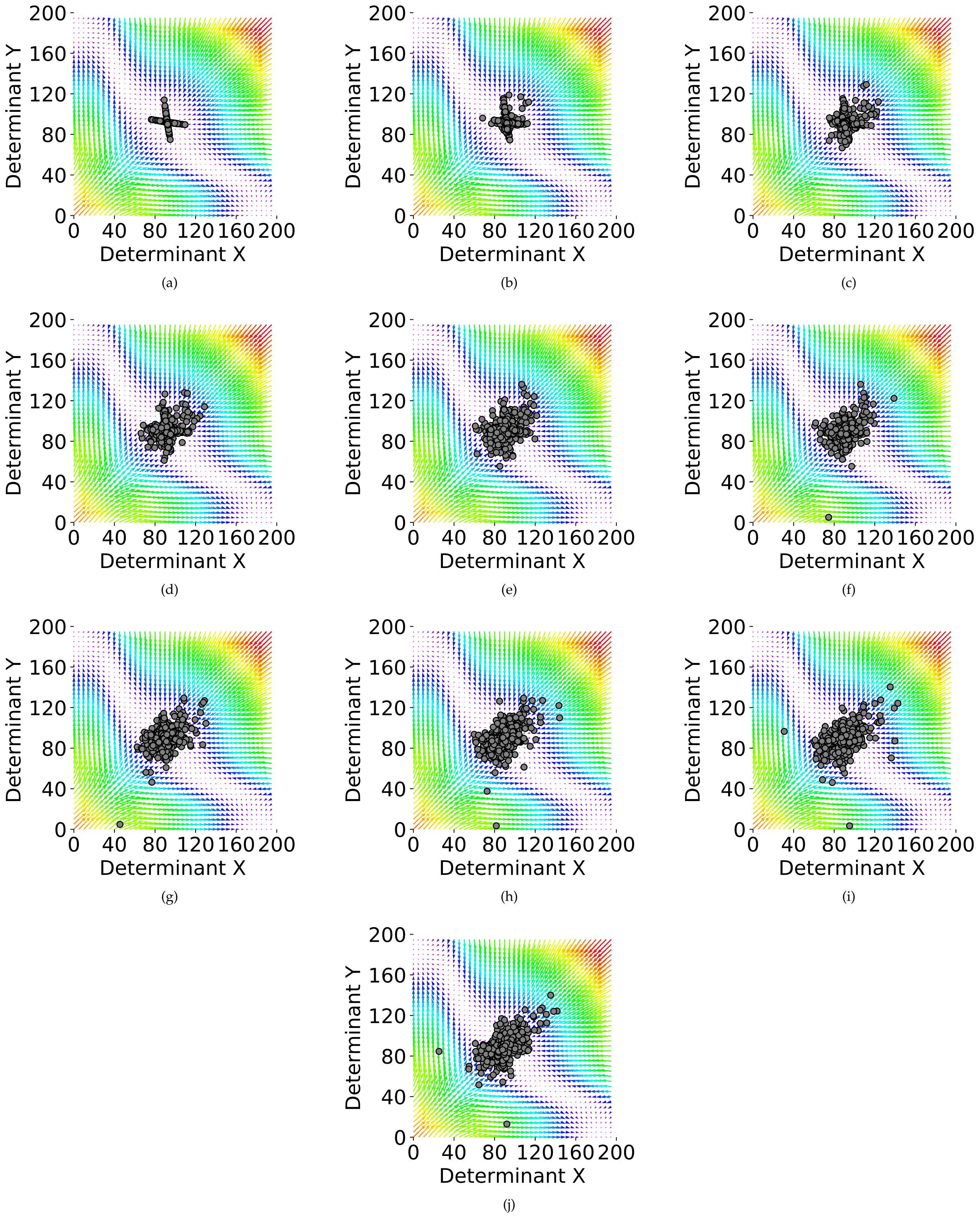


**Fig. S5. Ten phases of the simulation with the probability value of $p = 0.90$, and $\lambda = 5$. The internal regulatory networks of cells are assumed to be six-element switches.** (a-j) Phases 1 to 10 of the simulations. In each one of the plots, each circle represents the middle attractor of one of the cells in the population, with the representative cell being the one which produces the highest proportion of stem daughter cells at the end of each phase.

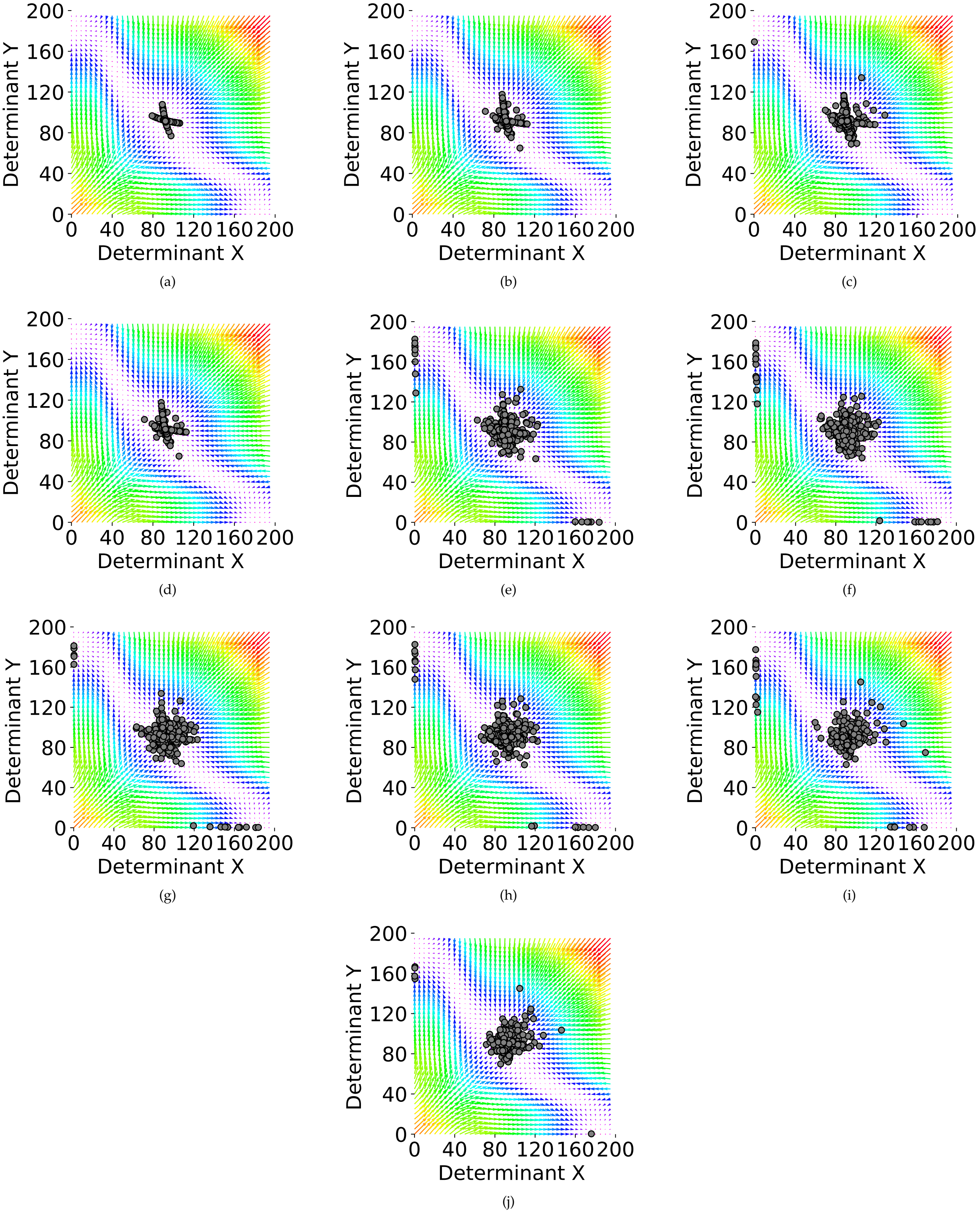


**Fig. S6. Ten phases of the simulation with the probability value of $p = 0.95$, and $\lambda = 5$. The internal regulatory networks of cells are assumed to be two-element switches.** (a-j) Phases 1 to 10 of the simulations. In each one of the plots, each circle represents the middle attractor of one of the cells in the population, with the representative cell being the one which produces the highest proportion of stem daughter cells at the end of each phase.

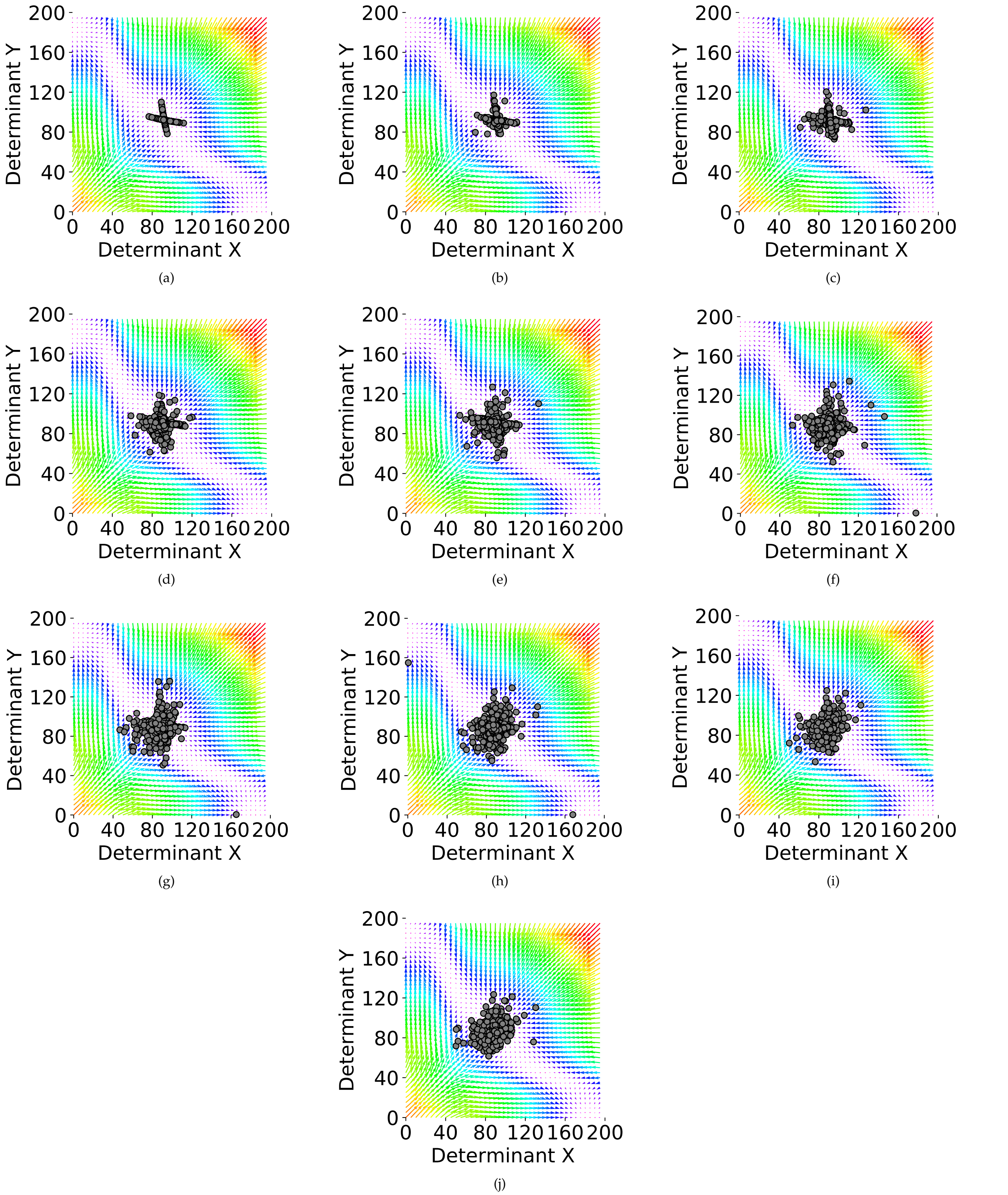


**Fig. S7. Ten phases of the simulation with the probability value of $p = 0.95$, and $\lambda = 5$. The internal regulatory networks of cells are assumed to be four-element switches.** (a-j) Phases 1 to 10 of the simulations. In each one of the plots, each circle represents the middle attractor of one of the cells in the population, with the representative cell being the one which produces the highest proportion of stem daughter cells at the end of each phase.

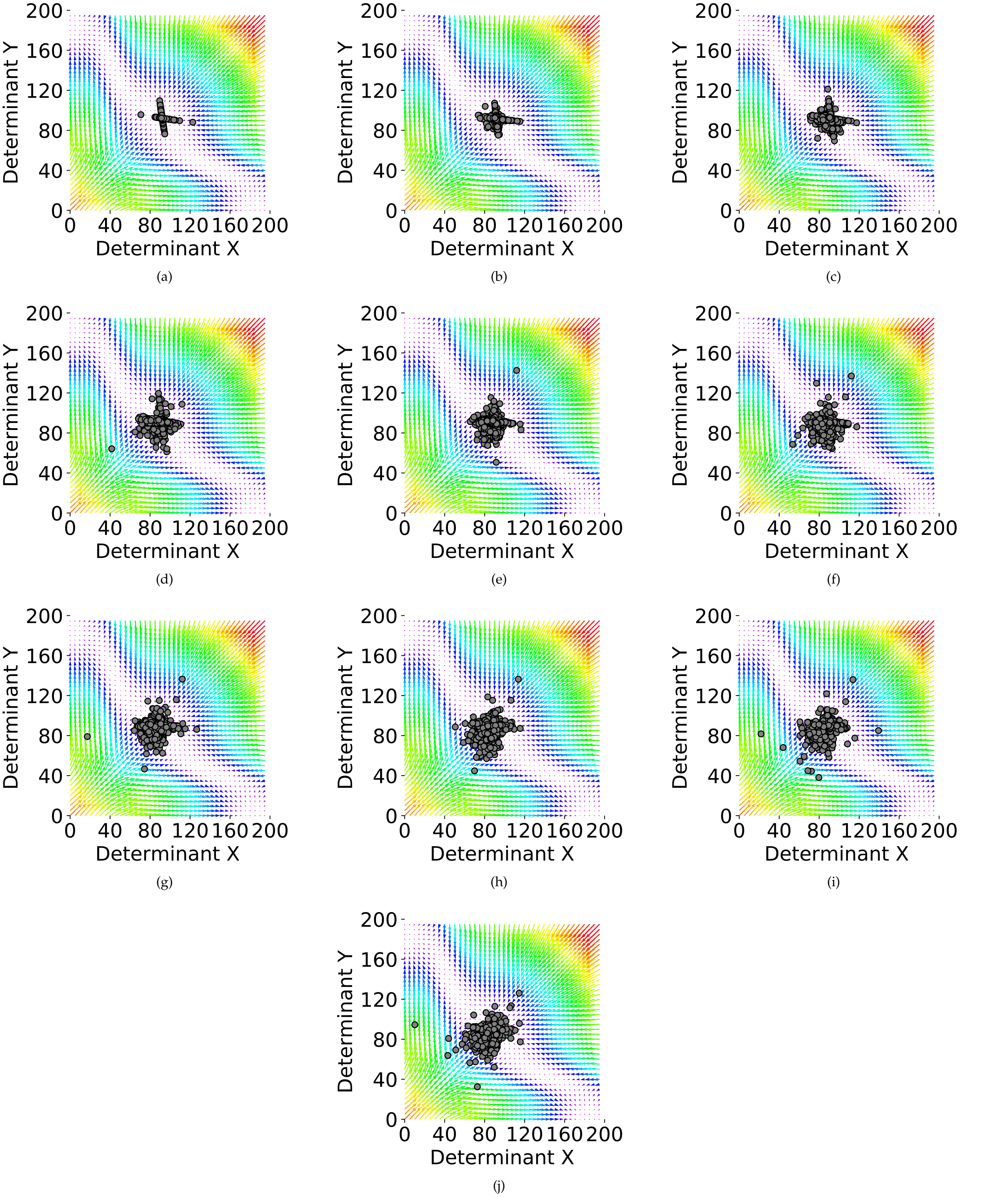


**Fig. S8. Ten phases of the simulation with the probability value of $p = 0.95$, and $\lambda = 5$. The internal regulatory networks of cells are assumed to be six-element switches.** (a-j) Phases 1 to 10 of the simulations. In each one of the plots, each circle represents the middle attractor of one of the cells in the population, with the representative cell being the one which produces the highest proportion of stem daughter cells at the end of each phase.

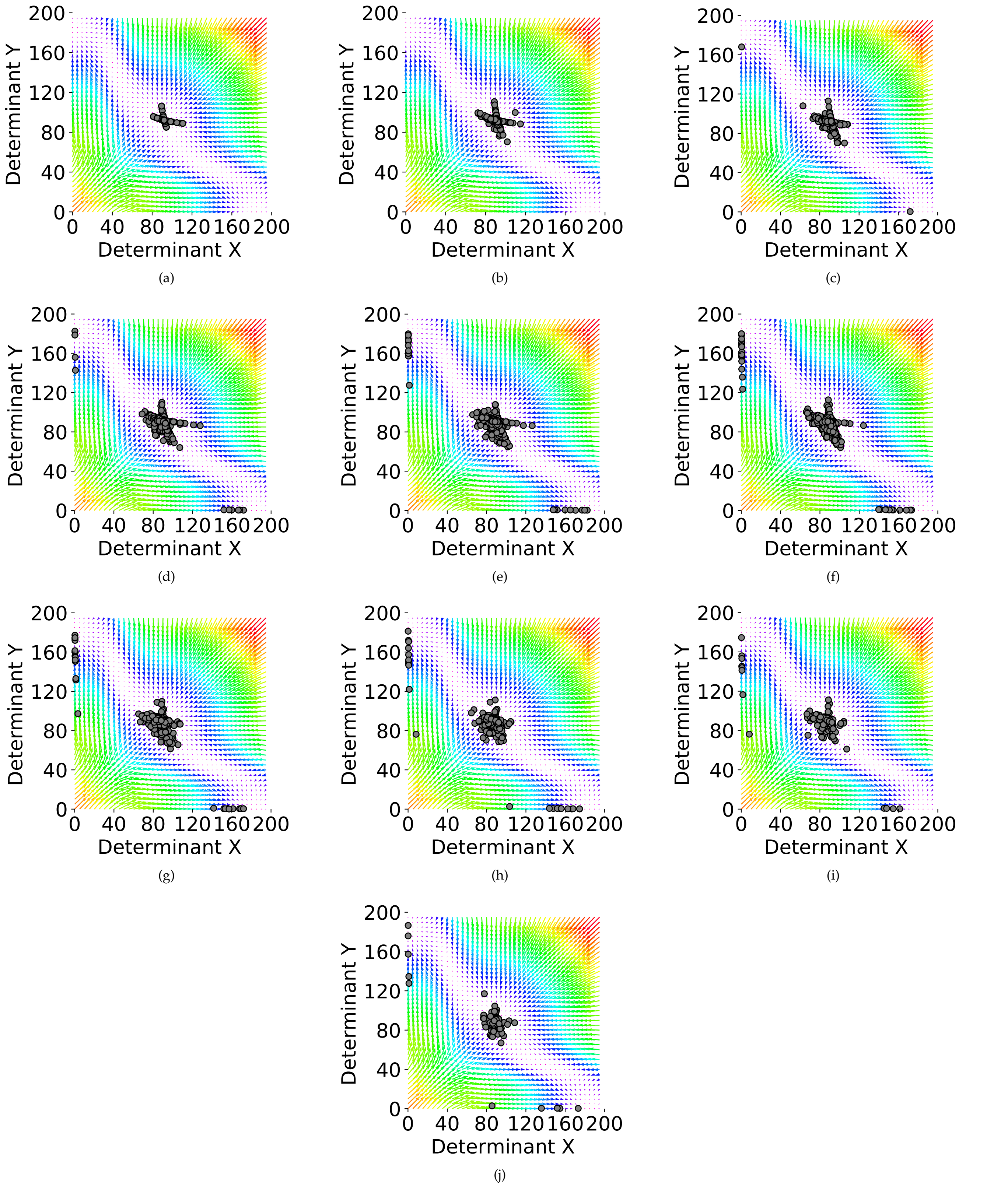


**Fig. S9. Ten phases of the simulation with the probability value of $p = 0.99$, and $\lambda = 5$. The internal regulatory networks of cells are assumed to be two-element switches.** (a-j) Phases 1 to 10 of the simulations. In each one of the plots, each circle represents the middle attractor of one of the cells in the population, with the representative cell being the one which produces the highest proportion of stem daughter cells at the end of each phase.

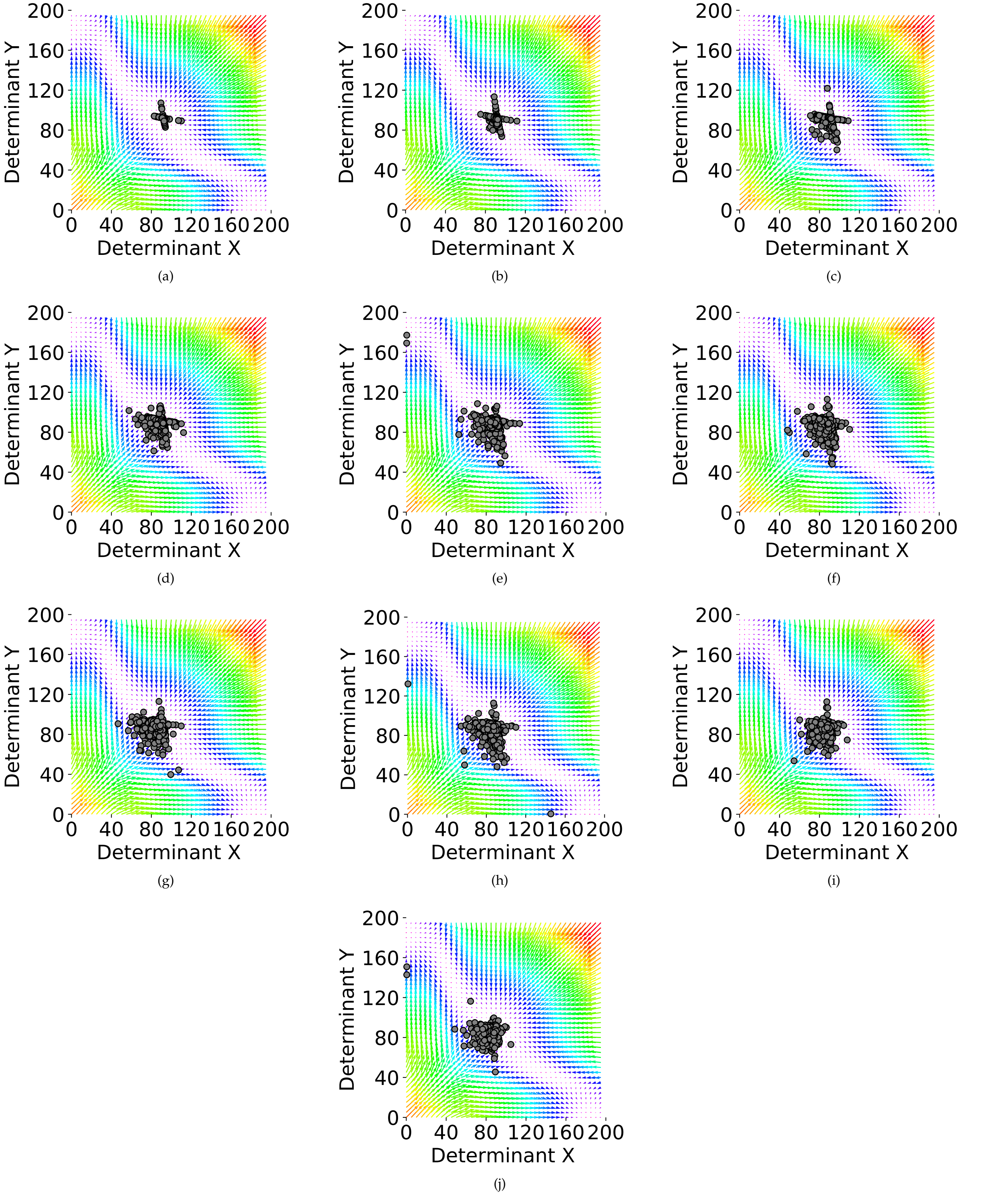


**Fig. S10. Ten phases of the simulation with the probability value of $p = 0.99$, and $\lambda = 5$. The internal regulatory networks of cells are assumed to be four-element switches.** (a-j) Phases 1 to 10 of the simulations. In each one of the plots, each circle represents the middle attractor of one of the cells in the population, with the representative cell being the one which produces the highest proportion of stem daughter cells at the end of each phase.

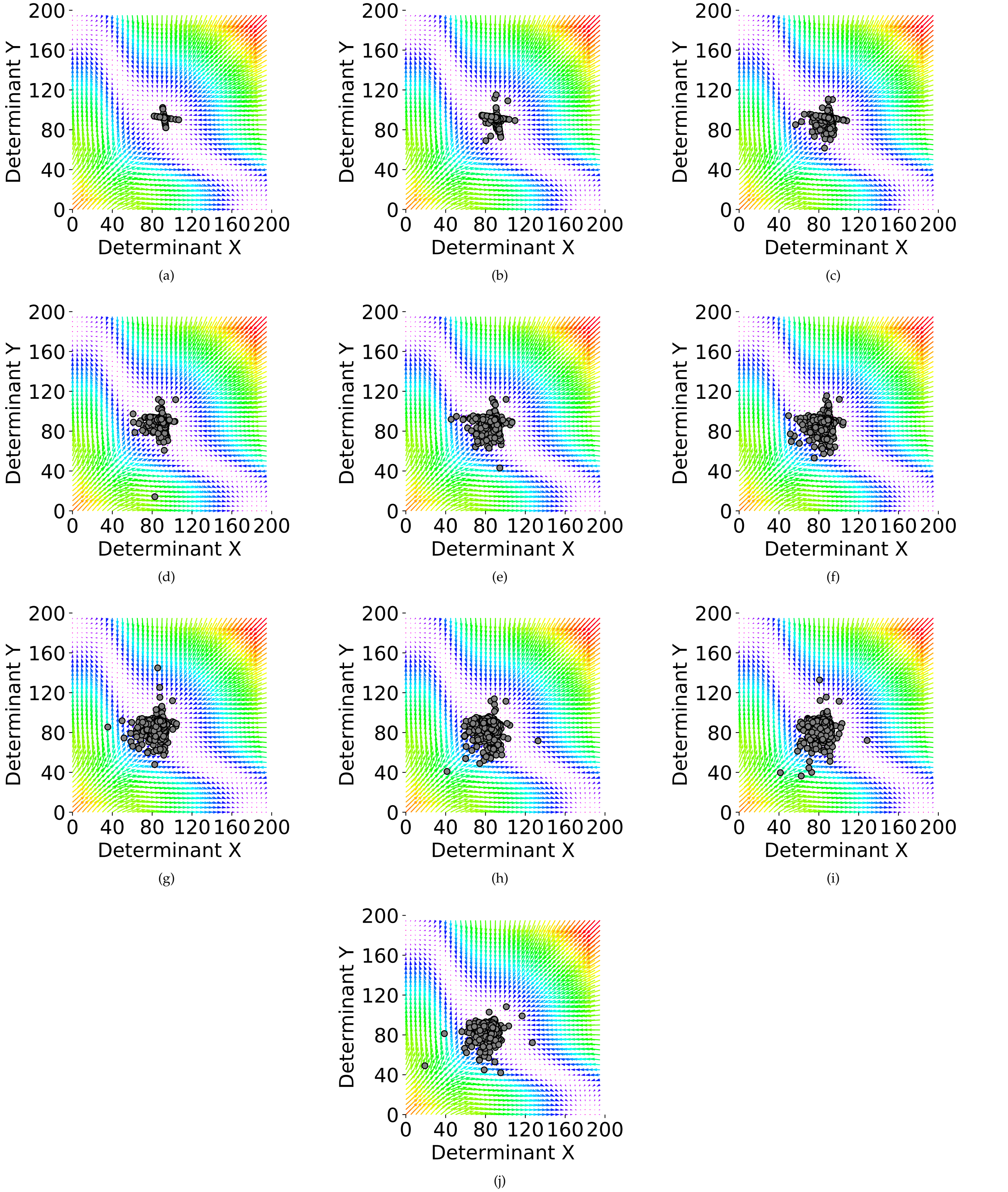


**Fig. S11. Ten phases of the simulation with the probability value of $p = 0.99$, and $\lambda = 5$. The internal regulatory networks of cells are assumed to be six-element switches.** (a-j) Phases 1 to 10 of the simulations. In each one of the plots, each circle represents the middle attractor of one of the cells in the population, with the representative cell being the one which produces the highest proportion of stem daughter cells at the end of each phase.

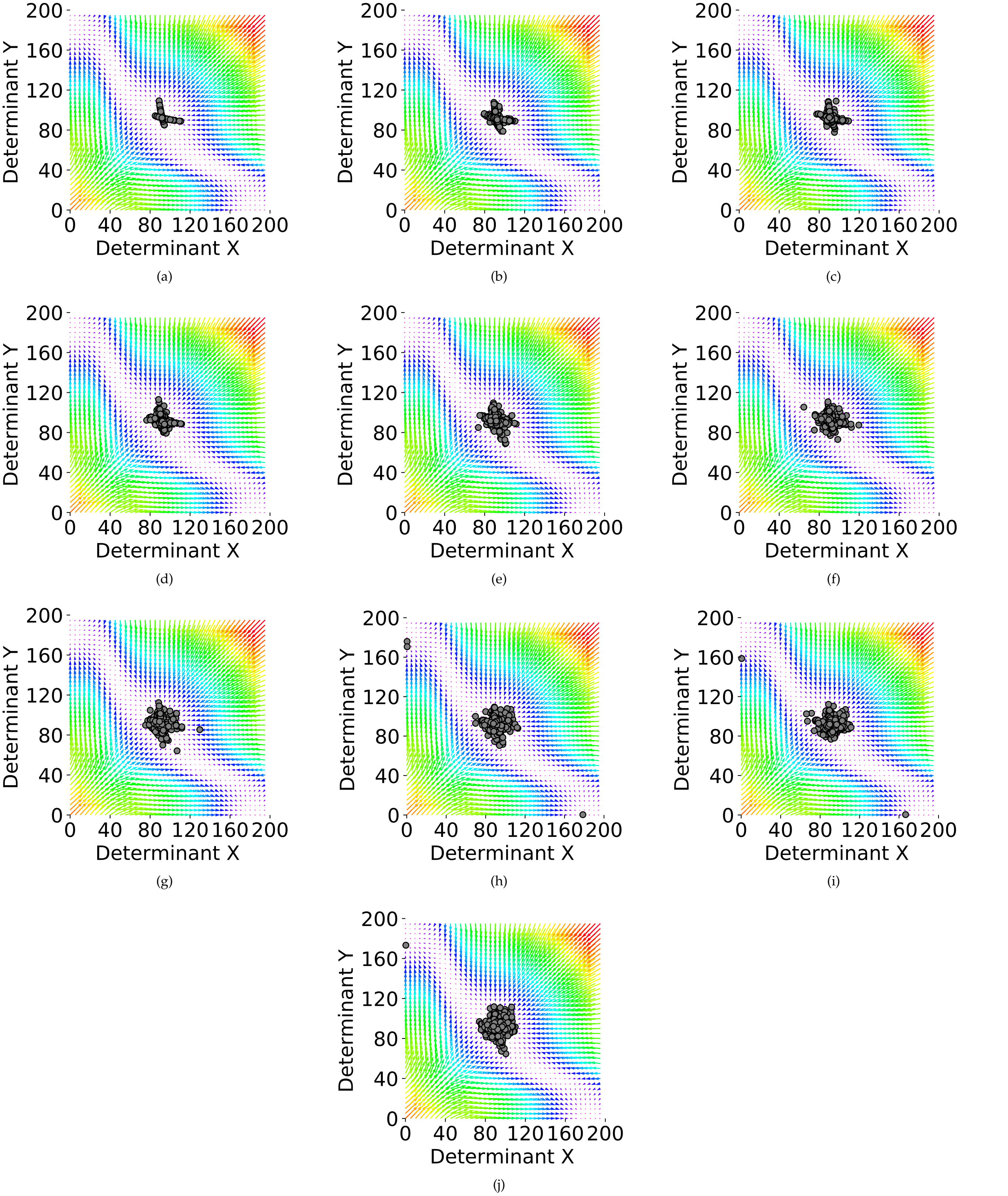


**Fig. S12. Ten phases of the simulation with the probability value of $p = 0.95$, and $\lambda = 2$. The internal regulatory networks of cells are assumed to be two-element switches.** (a-j) Phases 1 to 10 of the simulations. In each one of the plots, each circle represents the middle attractor of one of the cells in the population, with the representative cell being the one which produces the highest proportion of stem daughter cells at the end of each phase.

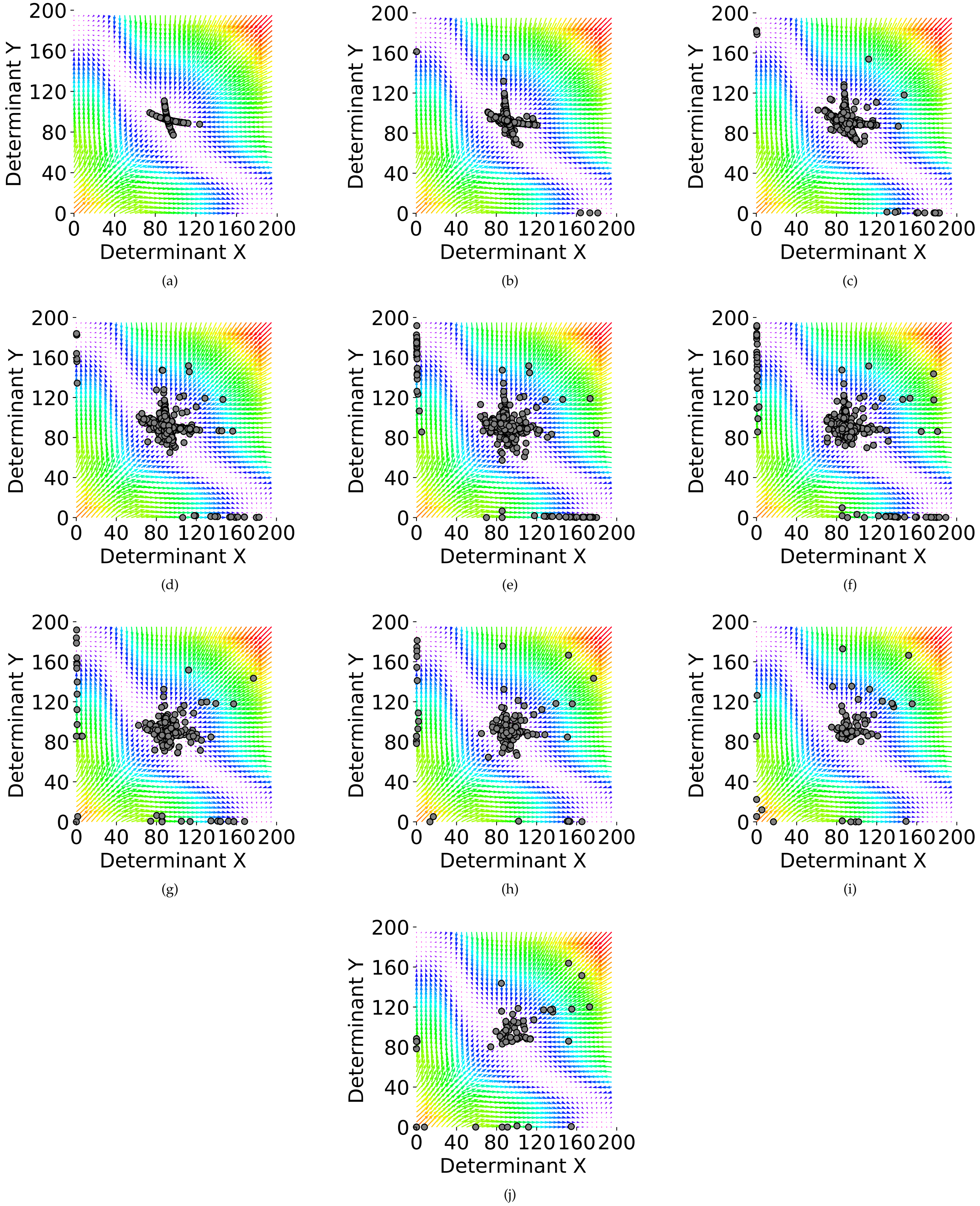


**Fig. S13. Ten phases of the simulation with the probability value of $p = 0.95$, and $\lambda = 10$. The internal regulatory networks of cells are assumed to be two-element switches.** (a-j) Phases 1 to 10 of the simulations. In each one of the plots, each circle represents the middle attractor of one of the cells in the population, with the representative cell being the one which produces the highest proportion of stem daughter cells at the end of each phase.
